# Symbiotic yeasts of a bark beetle transform major tree defenses into beetle protectants

**DOI:** 10.64898/2026.07.04.736475

**Authors:** Ana Patricia Baños-Quintana, Leandro Manuel Santiago-Padilla, Michael Reichelt, Ruo Sun, Martin Kaltenpoth, Jonathan Gershenzon, Maximilian Lehenberger

## Abstract

The Eurasian spruce bark beetle *Ips typographus*, a major forest pest on Norway spruce (*Picea abies*), forms intimate associations with several types of microbial symbionts. While previous research has focused primarily on filamentous fungi, yeasts have remained largely unexplored. Here, we show that yeasts associated with *I. typographus* may contribute to host tree colonization by providing defensive benefits. Dominant yeasts (*Yamadazyma*, *Kuraishia*, *Nakazawaea,* and *Wickerhamomyces*), which are phylogenetically related to other insect-associated Saccharomycotina, significantly attract adult beetles. Moreover, several yeasts inhibit the growth of the pathogenic fungus *Trichoderma harzianum in vitro*, and beetle eggs benefit from the presence of *Kuraishia capsulata* by reduced fungal infection under semi-natural conditions. Strikingly, these effects are mediated by the yeasts’ transformation of the tree’s defensive stilbene glycosides into antimicrobial aglycones and phenolic acids that accumulate in beetle galleries. These findings reveal a previously unrecognized role of symbiotic yeasts in converting spruce defensive stilbene glycosides into antimicrobial aglycones and oxidative cleavage products that accumulate in beetle galleries, and might contribute to the survival of their bark beetle host.

## Introduction

Bark beetles (Curculionidae: Scolytinae) are the main biotic driver of disturbances in temperate forests of the Northern Hemisphere (1,2). With annual losses of up to 70 Mm³ of timber in Europe alone (1), the Eurasian spruce bark beetle (*Ips typographus*) is one of the most devastating forest pests in the Palearctic. Its ecological success is astonishing, given that its host tree, Norway spruce (Pinaceae: *Picea abies*), possesses very effective constitutive and inducible defense mechanisms against herbivore attacks (3). Spruce trees have thick, suberized cork that serves as an outermost protective layer to the stems, as well lignin-rich stone cells scattered throughout the phloem which hinder the advance of herbivores (3). Aside from these physical barriers, Norway spruce also produces constitutive chemical defenses: polyphenolic parenchyma cells containing stilbene glycosides and flavonoids are abundant in the phloem, and resin ducts store toxic and viscous terpenoid mixes in the xylem and phloem (3). Upon bark beetle attack, the tree’s inducible chemical defenses are activated: the *de novo* production of terpenoids and phenolic compounds is upregulated, creating an even more hostile environment for invaders (4,5). Despite this multi-layered defense system, *I. typographus* are able to colonize and kill large numbers of *P. abies* during eruptive beetle population phases. While several climatic and anthropogenic factors are known to favor bark beetle outbreaks, the biological aspects behind the insects’ destructive capabilities are less understood (6).

Symbioses with microorganisms are widespread across insect orders, and they are one of the evolutionary mechanisms underlying insects’ adaptations to challenging environments and nutrient-poor diets (7). Bark beetles form partnerships with filamentous fungi of diverse taxonomic groups, and these associations vary in their degree of specialization (8). In some species, beetles have even evolved dedicated structures (*i.e.*, mycetangia) to transport their fungal ectosymbionts and vector them into the host trees during colonization (9). In the case of *I. typographus,* the beetle associates with several Sordariomycetes (Ascomycota) that utilize the tree’s phenolic compounds as carbon sources (10) and transform resin monoterpenes into volatile cues that aid the beetles in locating them (11). Thus, it is likely that one of their main roles consists of metabolizing spruce defense compounds during the attack (8). Further, it has been hypothesized that these fungi may serve as an additional nutritional source for the beetle and protect it from pathogenic microorganisms during host tree invasion, but these roles have not been experimentally demonstrated (8). Other members of the bark beetle microbiota are underrepresented in the literature and less is known about their contributions to the insect’s success (12,13).

The association of bark beetles with yeasts has been known since the 1930s (14), yet, their ecology and a potential functional role within the beetle-microbe-tree system remain poorly investigated (12). A consistent group of yeasts is found in the gut and gallery environment of *I. typographus* throughout its life stages and across different geographic locations in Eurasia: taxa from the genera *Wickerhamomyces*, *Kuraishia*, *Nakazawaea, Yamadazyma, Ogataea, Cryptococcus, and Cyberlindnera* are among the most abundant members of its fungal communities (15–18). Female beetles introduce yeasts into the phloem during tree colonization, making them available to their offspring in the oviposition sites (18). This vertical transmission mode, along with the ubiquitous presence of yeasts belonging to these genera in other *Ips* (19) and *Dendroctonus* (20,21) bark beetles, suggest that these microbes are important for Scolytines (12). *I. typographus*-associated yeasts can convert *cis*-verbenol and *trans*-verbenol, major components of the beetle’s aggregation pheromone, into verbenone (22), an anti-aggregation pheromone that the insect utilizes to avoid intraspecific competition when choosing a new host tree (23). Aside from emitting volatile signaling compounds, several of these yeasts possess complete metabolic pathways for essential amino acid and vitamin B6 biosynthesis, as well as large numbers of genes related to detoxification processes (24), but their relevance for bark beetles’ survival remains unknown.

In this study, we investigated the roles of the yeast communities in the successful colonization of *I. typographus* in its host tree, Norway spruce. We hypothesized that yeasts are involved in metabolizing some of the tree’s major defense compounds. Chemical analyses coupled with *in vitro-* and *in vivo*-bioassays reveal that the yeasts transform the tree’s defenses into compounds that may be used for protecting the beetle eggs against pathogens. These findings highlight a previously underappreciated role of yeast communities in the life history of bark beetles.

## Results

### *Ips typographus* harbors a characteristic community of yeasts and is attracted to its members

To study the yeast community associated with *I. typographus,* we generated an isolate collection for phylogenetic analysis, behavioral assays and further biochemical characterization. A total of 76 yeast isolates were obtained from the surfaces, heads and abdomens of adult *I. typographus* by culture-dependent methods and identified by sequencing the gene encoding the large subunit (28S) of the rRNA (Suppl. Table S1, Suppl. Fig. S1). The most abundant taxa, *Yamadazyma scolyti, Y. tenuis, Wickerhamomyces bisporus, Kuraishia capsulata, K. molischiana,* and *Nakazawaea holstii*, were selected for subsequent analysis. The phylogenetic placement of these species showed that they are related to yeasts that were isolated from insects, their habitats, or decaying wood (Fig. 1). Olfaction-guided choice assays with *in vitro* cultures (Fig. 2a) showed that adult *I. typographus* were significantly attracted to their main yeast associates versus uninfested medium (exact binomial test *p <* 0.05, n = 40, Fig. 2b), except for *W. bisporus* (*p =* 0.4296). When presented with the yeast *Danielozyma ontarioensis*, a symbiont of another bark beetle *Polygraphus poligraphus* (25), the beetles did not prefer this isolate over the uninfested control (*p =* 0.6358). Further, the insects showed a strong aversion towards the pathogenic fungus *Trichoderma harzianum* (*p =* 1.383×10^-06^), which co-occurs in bark beetle galleries (11). Our culture-dependent characterization of the yeast community complements previous culture-independent analyses that found the same taxa associated with *I. typographus* in different geographic locations and across beetle life stages (15–18). Taken together, the consistency of these associations, the close phylogenetic relationship of the isolates to other insect-associated yeasts, and the attractive effect of yeast volatiles on adult beetles indicate a stable symbiosis that may be beneficial to both insect and yeast partners.

**Figure 1.**
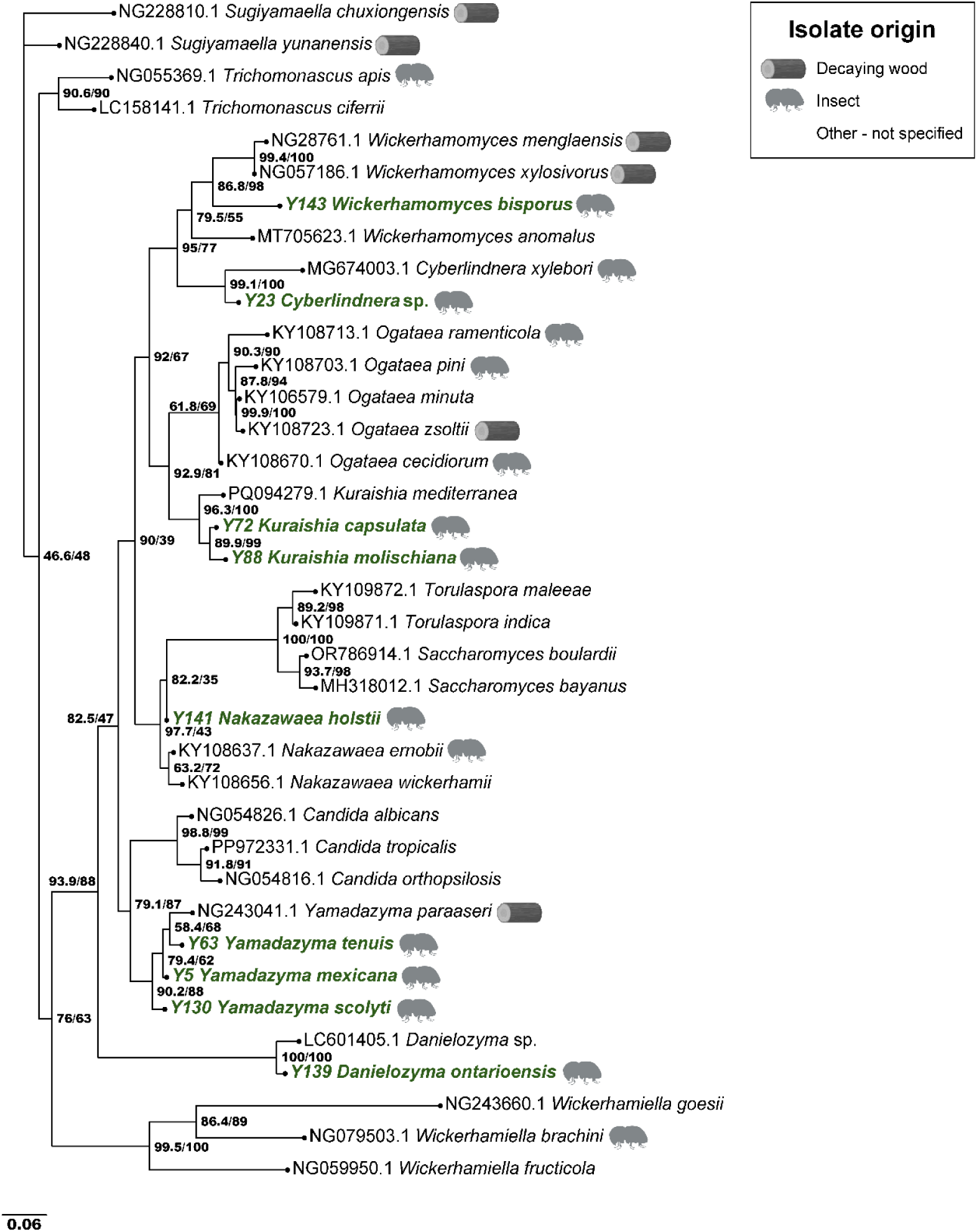
Phylogenetic placement of *Ips typographus*-associated yeasts. Maximum likelihood phylogeny of the yeast species associated with the bark beetle *I. typographus* investigated in this study (highlighted in green) based on fungal LSU sequences. The sequences of nine yeast species were generated in this study, while the remaining 28 sequences were retrieved from the NCBI GenBank database. Shimodaira-Hasegawa approximate likelihood ratio test (SH-aLRT) support values (left numbers) and ultrafast bootstrap support values (right numbers) are provided in the phylogenetic placement. The strain identity (NCBI accession number or isolate ID generated in this study) is indicated on the left of the species name. In cases where the isolates were obtained from insect hosts or from decaying wood, the source is indicated with a pictogram.

**Figure 2.**
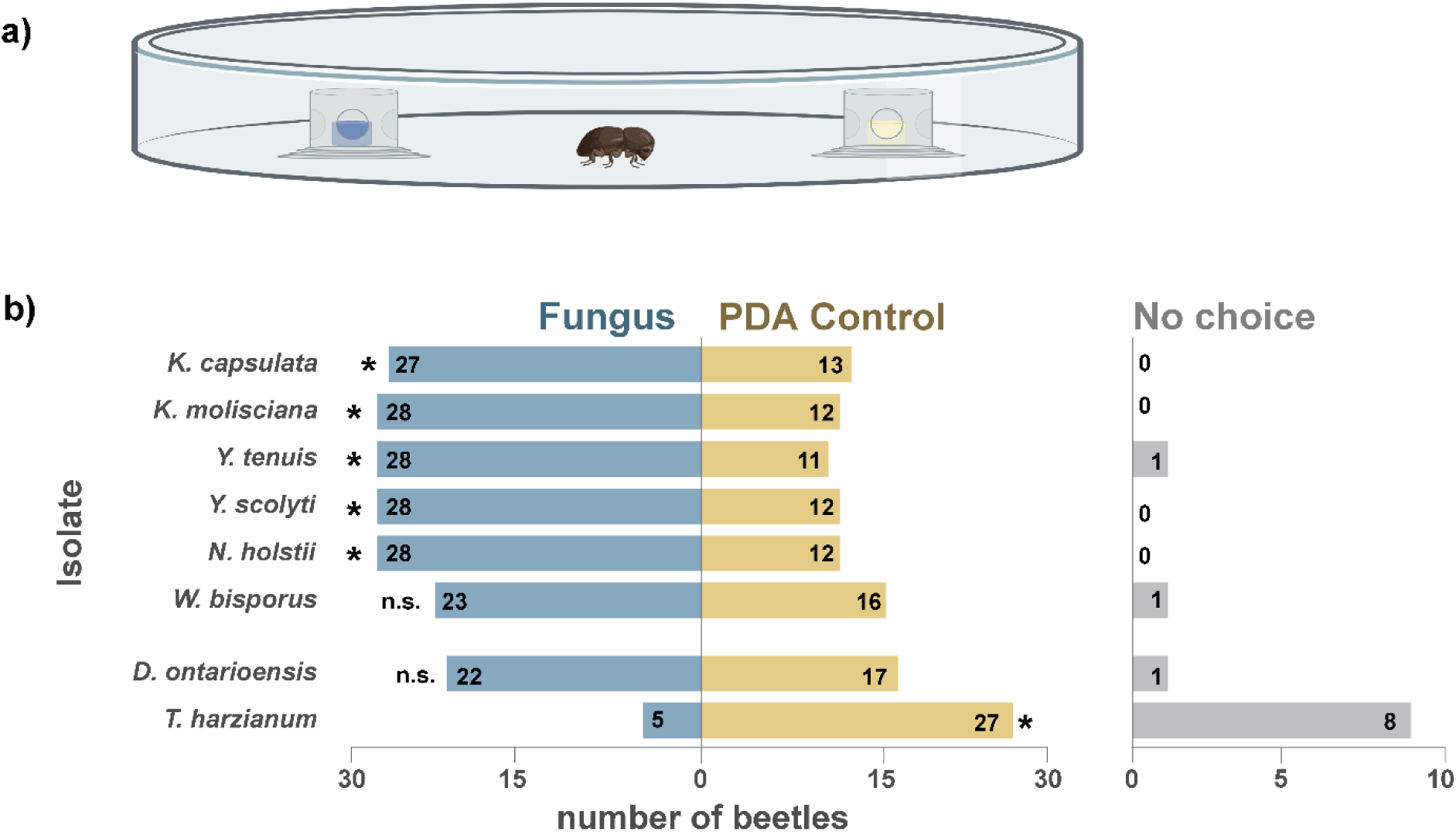
Olfaction-guided preference assays. **a)** Adult *I. typographus* were placed in an arena that consisted of a large Petri dish containing two equidistant cups provisioned with either a fungal isolate or a PDA control. Beetles chose between samples guided only by olfactory cues. **b)** Number of beetles that chose a fungal isolate (blue) or the PDA medium control (yellow). The beetles that did not enter any of the sample-containing cups after 6 hours were recorded as “no choice” (gray). Statistically significant differences are indicated by an asterisk (exact binomial test, *p* < 0.05, n. s. = not significant, n = 40).

### Yeasts protect beetle eggs from a pathogenic fungus

Bark beetles and their symbionts encounter a microbe-rich environment upon arriving at a new spruce host tree. However, during the excavation of the maternal and larval galleries, the composition of the fungal and bacterial communities shifts and becomes dominated by a subset of taxa associated with the bark beetles. Yeasts are early microbial colonizers of the galleries and remain some of the most abundant fungi throughout the development of the beetles (15,18). We hypothesized that yeasts can regulate the growth of other fungi in the galleries (12) by creating unfavorable growth conditions for non-beneficial saprophytes and pathogens. To test this, we carried out *in vitro* assays where the pathogenic fungus *Trichoderma harzianum* was confronted with each of the main bark beetle-associated yeasts (Fig. 3a). When co-cultivated on potato dextrose agar (PDA), *T. harzianum* colonized the full plate, and no inhibition zones were formed around any of the yeast isolates. However, when *T. harzianum* and the yeasts were co-cultivated on the more ecologically relevant spruce phloem agar (SPA), *K. capsulata, K. molischiana* and *N. holstii* inhibited the growth of *T. harzianum* (Fig. 3b). When we repeated these confrontation assays with two major bark beetle ectosymbionts, *G. penicillata* and *E. polonica*, and three of the yeasts, the growth of *G. penicillata* remained unaffected by the presence of yeasts, while *E. polonica* displayed delayed growth but was not significantly inhibited (Suppl. Fig. S2).

**Figure 3.**
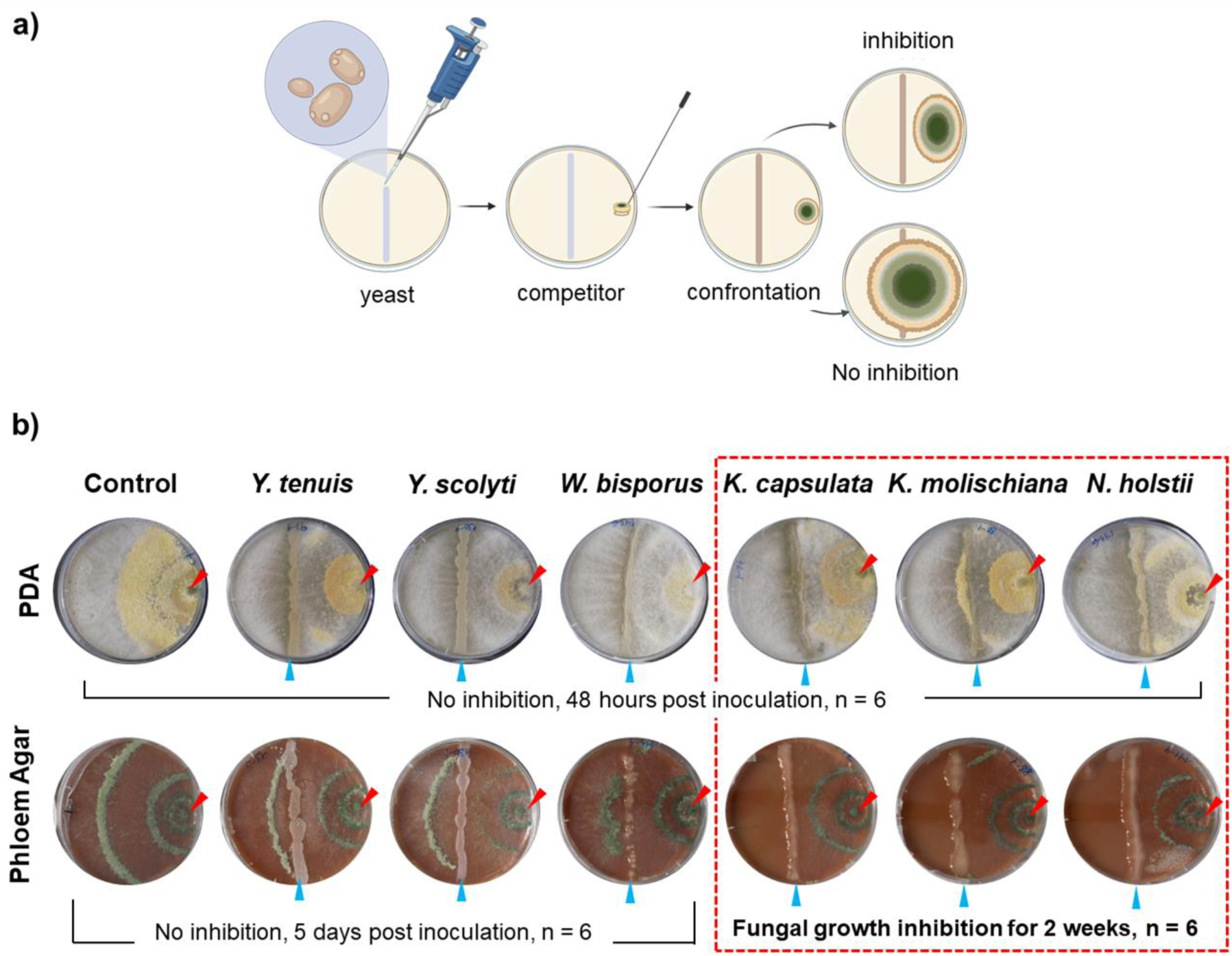
Confrontation assays: yeasts against the pathogen *Trichoderma harzianum.* **a)** Schematic view of the assay setup. A yeast suspension was pipetted on an agar plate forming a straight line crossing the center. A 4 mm plug of *T. harzianum* culture was placed at one of the ends of the plate. The fungi were co-cultivated for up to two weeks and pictures were taken every day to record fungal growth. **b)** Confrontation assay results on potato dextrose agar medium (PDA, top row) and rich spruce phloem agar medium (SPA amended with 1% PDA, bottom row). A blue arrow indicates the inoculation line of the yeast and a red arrow the placement of *T. harzianum* plugs. Yeast species names are shown at the top; those that inhibited *T. harzianum* growth when growing on SPA (and not when growing on PDA) are enclosed in a red dotted box. *T. harzianum* inoculated in the absence of yeasts was used as a control, n = 6 plates.

Female *I. typographus* lay their eggs in individual niches that they carve out on the sides of the maternal gallery and cover them with a protective plug made from masticated phloem. The females inoculate the plug with their associated yeast and bacteria (18). To test whether the antifungal effect displayed by the yeasts is ecologically relevant for the beetle’s early developmental stages, we generated yeast-free eggs by surface-sterilization and exposed them to *T. harzianum* under semi-natural conditions (26). Briefly, surface-sterilized eggs were randomly assigned to either sterile or *Kuraishia capsulata*-inoculated artificial phloem plugs, and then challenged with a suspension of *T. harzianum* spores (Suppl. Fig. S3). Untreated eggs placed on sterile artificial phloem plugs were used as a negative control. The *T. harzianum* infection probability was significantly lower when the yeast *K. capsulata* was inoculated into the plugs at concentrations comparable to those found under field conditions (18) (Fig. 4, Log-rank test X^2^ = 11.5, *p =* 0.003; Cox proportional hazards regression *p =* 0.002 for the presence of *K. capsulata*; n = 18). The egg’s native microbiota alone (control treatment) was insufficient to confer protection against the pathogen *T. harzianum*, which highlights the importance of yeast inoculation by female beetles into the phloem plugs during oviposition.

**Figure 4.**
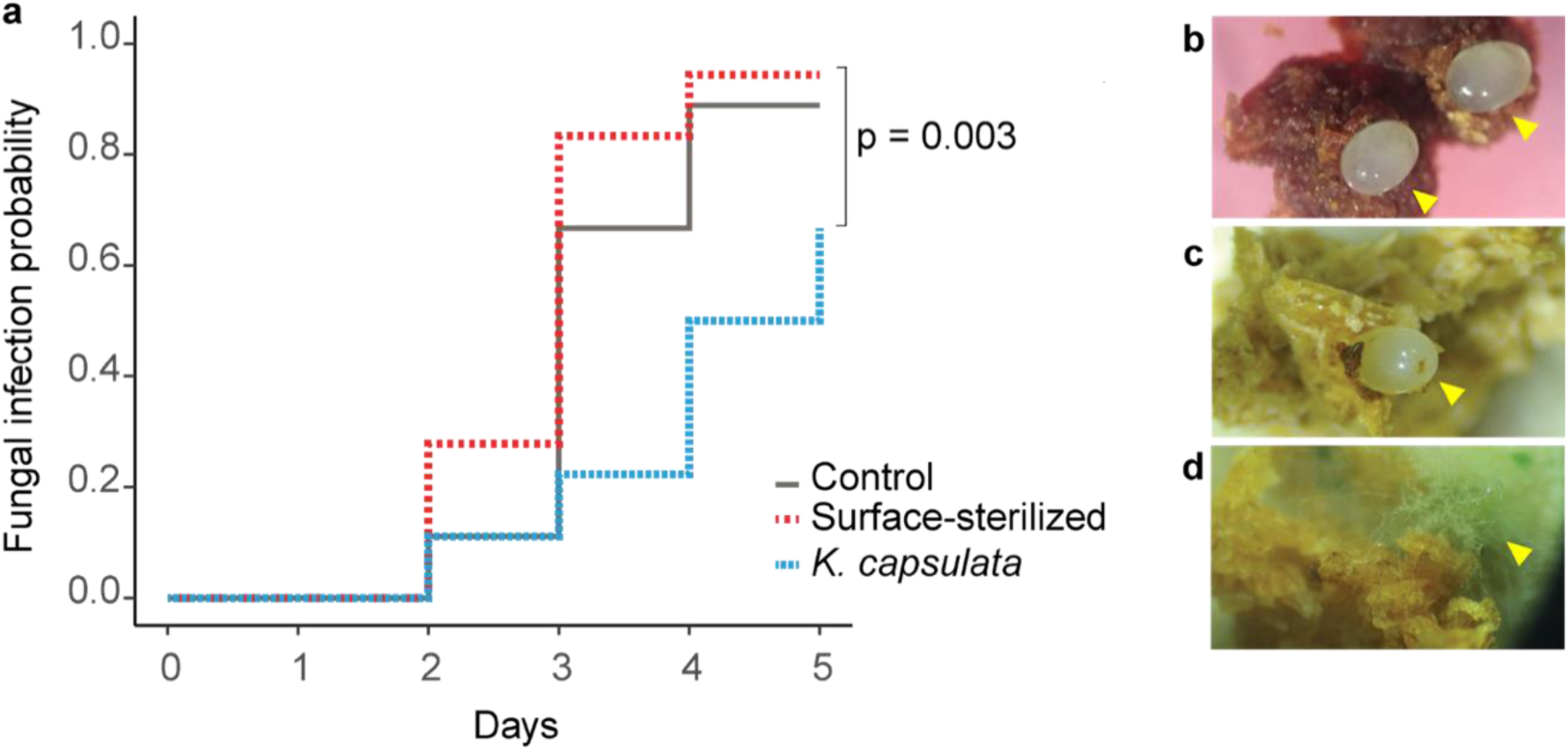
Egg infection bioassay. **a)** Kaplan-Meier estimate of *T. harzianum* infection probability after 5 days of exposure to a spore suspension of this pathogen. Surface-sterilized *I. typographus* eggs were randomly assigned to sterile artificial phloem plugs (‘Surface-sterilized’ treatment, red) or phloem plugs inoculated with the yeast *Kuraishia capsulata* (‘*K. capsulata*’ treatment, blue). Untreated eggs with their native microbiota were placed on sterile artificial phloem plugs and used as a control (‘Control’ treatment, gray). Log-rank test X^2^ = 11.5, *p* = 0.003; Cox proportional hazards regression *p* = 0.002 for the presence of *K. capsulata* compared to the other treatments; n =18. **b)** Eggs on natural phloem plugs produced by female *I. typographus* in the oviposition sites. **c)** Uninfected and **d)** *Trichoderma-*infected egg on an artificial phloem plug, five days post exposure to the pathogen’s spores. Yellow arrows indicate the position of the eggs.

### Yeasts metabolize a group of major defense compounds in spruce phloem

Since the antifungal effect of the cultivated yeasts was only observed in the presence of spruce phloem, we hypothesized that yeasts utilize compounds present in Norway spruce or their metabolites to regulate the growth of filamentous fungi in the galleries. The stilbene glycosides piceid, astringin and isorhapontin are among the most abundant defense compounds in *P. abies* phloem (27). LC-MS/MS analysis of freeze-dried spruce phloem agar revealed that the content of the three stilbene glycosides was significantly reduced after five days of incubation with the individual yeasts, reaching near-zero concentrations in all cases (Fig. 5). This also occurred in fresh spruce phloem agar (Suppl. Fig. S4) and fresh yeast biomass (Suppl. Fig. S5). Conversely, we observed a significant increase of the corresponding stilbene aglycones resveratrol, piceatannol and isorhapontigenin in the phloem agar (Fig. 5; Suppl. Fig. S4) and the yeast biomass (Suppl. Fig. S5) of all isolates except for *K. capsulata*. This yeast enriched both spruce phloem and its own biomass with the phenolic acids 4-hydroxybenzoic acid, protocatechuic acid and vanillic acid (Fig. 5, Suppl. Figs S4-S5), likely derived by oxidative cleavage of the corresponding stilbene aglycones (28). These results suggest that *I. typographus-*associated yeasts metabolize the stilbenes present in spruce phloem.

**Figure 5.**
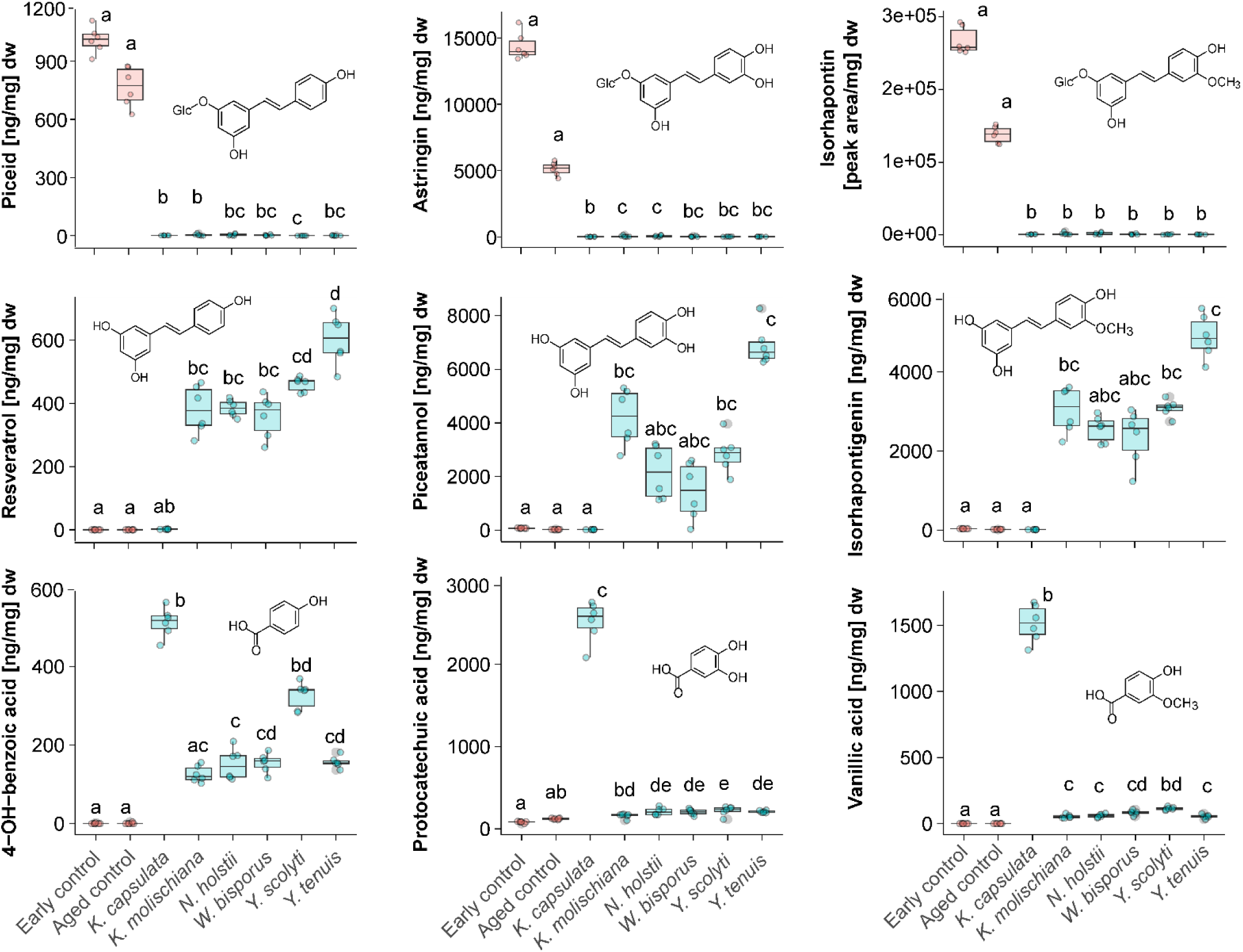
Stilbene and phenolic acid content in spruce phloem agar after yeast growth. The concentration of stilbene glycosides (top row), their aglycones (middle row) and the corresponding phenolic acids (bottom row) were measured in spruce phloem agar medium five days after inoculation with *I. typographus-*associated yeasts. Non-inoculated spruce phloem agar was sampled at the start of the experiment (‘Early control’) and after five days of incubation (‘Aged control’) to be used as a control. Concentrations are shown in nanograms per milligram of dry weight except for isorhapontin, where the peak area was used instead since no standard was available for calibration. Different letters indicate significant differences among groups (Kruskal-Wallis rank-sum test followed by Dunn’s post-hoc test, *p* < 0.05, n = 6 plates).

To assess whether our *in vitro* findings reflect the chemical changes that occur in a natural setting, we sampled bark beetle galleries at multiple time points after beetle infestation and measured the concentrations of stilbene glycosides, stilbene aglycones and phenolic acids. We observed a strikingly similar pattern, with high stilbene glycoside contents in non-excavated phloem adjacent to beetle galleries and near-zero concentrations of these compounds in excavated galleries collected at the same time point (Fig. 6; Suppl. Fig. S6). Further, the stilbene aglycone content was significantly higher in early larval galleries (Fig. 6), recently excavated maternal galleries, egg plugs, and the frass collected from pupal chambers than in non-excavated adjacent phloem (Suppl. Fig. S6). Phenolic acids were also detected in the gallery environment, especially in freshly excavated maternal galleries (Suppl. Fig. S6). The concentrations of both the stilbene aglycones and phenolic acids decayed over time, which may be partially explained by the ecological succession of microorganisms within the galleries. Highly specialized, later-colonizing fungal species may metabolize stilbene aglycones or even use them as a carbon source, as has been shown for some of the filamentous symbionts of *Ips typographus* (10,27,29). As for phenolic acids, an *in vitro* assay with a panel of bark and ambrosia beetle-associated filamentous fungi showed that most of them are capable of metabolizing these molecules after 12 days (Suppl. Fig. 7).

**Figure 6.**
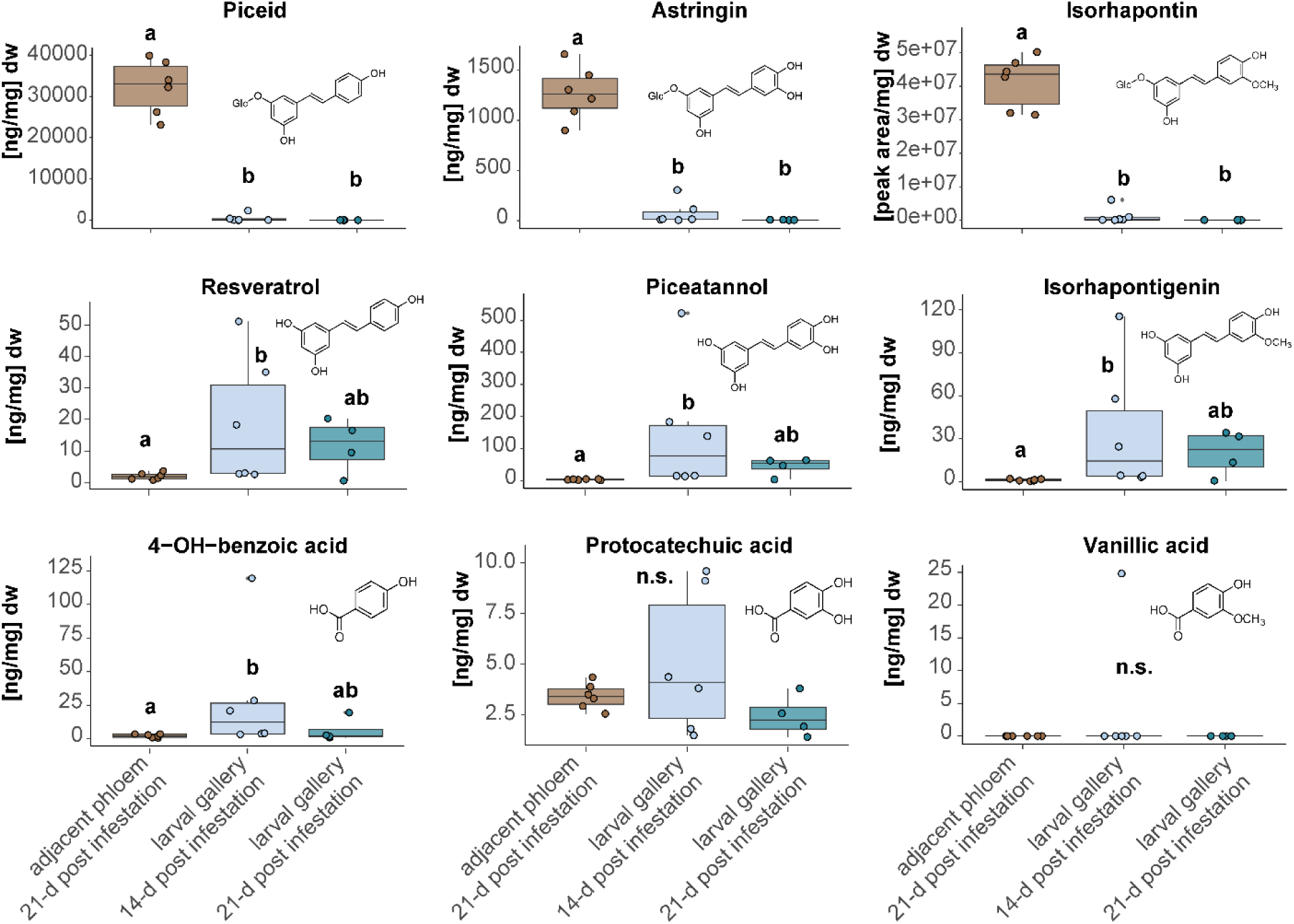
Stilbene and phenolic acid content in the larval galleries and the adjacent phloem at different timepoints after bark beetle infestation. The concentration of stilbene glycosides (top row), their aglycones (middle row) and the corresponding phenolic acids (bottom row) were measured in the larval galleries 14 and 21 days after exposing spruce logs to laboratory-reared *I. typographus* adults. The unattacked phloem adjacent to the galleries was used as a control. Concentrations are shown in nanograms per milligram of dry weight except for isorhapontin, where the peak area was used instead. Different letters indicate significant differences among groups (Kruskal-Wallis rank-sum test followed by a Tukey post-hoc test, *p* < 0.05, n = 4-6 galleries).

### Yeast stilbene metabolites inhibit the growth of a pathogenic fungus

As stilbene aglycones and phenolic acids were detected as major yeast-produced derivatives of the stilbene glycoside tree defenses (Fig. 5; Suppl. Figs. S4-S5), and these compounds are known to have antifungal properties (30,31), we evaluated their bioactivity in mixtures against bark beetle-associated fungi using an agar strip assay. The assay was set up by cutting two-centimeter strips out of the center of PDA plates and then refilling them with PDA containing either 2% DMSO (control), a stilbene aglycone mix, or a phenolic acid mix that emulated the concentrations produced by the yeasts in spruce phloem agar. A fungal inoculum was placed on one side of each plate and incubated until the mycelium of the control treatment reached the opposite edge of the plate (Fig. 7a). The stilbene aglycone mix containing resveratrol, piceatannol and isorhapontigenin showed a strong inhibitory effect on the pathogenic fungus *T. harzianum* and reduced the growth of the beetle symbiont *E. polonica* (Fig. 7b, Kruskal-Wallis rank-sum test followed by Dunn’s post-hoc test, *p <* 0.05). Interestingly, the beetle symbiont *G. penicillata* was not significantly inhibited by the stilbene aglycone mix (Fig. 7b, *p =* 0.24). The phenolic acid mix containing 4-hydroxybenzoic, vanillic, and protocatechuic acids did not inhibit the growth of any of the fungi (Fig. 7). These results are congruent with the observations from the co-cultivation assays of filamentous fungi and the yeasts *K. capsulata*, *K. molischiana*, and *N. holstii* (Fig. 3 and Suppl. Fig. S2), indicating that stilbene aglycones may form part of the yeasts’ chemical defenses against fungal competitors in the galleries.

**Figure 7.**
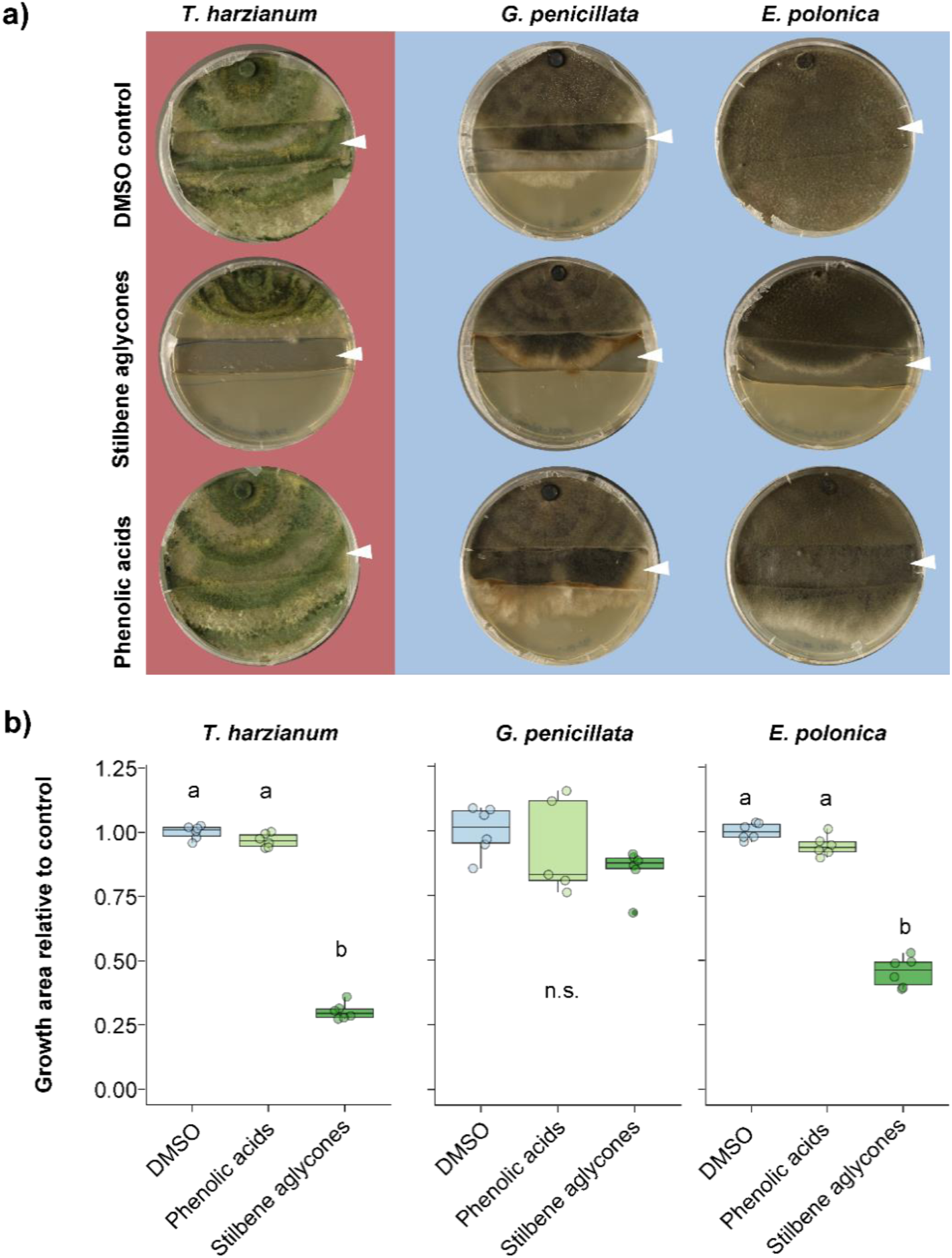
Agar strip assays with different stilbene compound mixes to assess anti-fungal activity. **a)** Two-centimeter-wide strips were removed from the middle of potato dextrose agar (PDA) plates to create a well. The well was subsequently filled with PDA amended with pure DMSO (2% DMSO, control treatment), a stilbene aglycone mix (piceatannol, resveratrol and isorhapontigenin at 500 μg/g medium each), or a phenolic acid mix (protocatechuic, vanillic, and 4-hydroxybenzoic acids at 500 μg/g medium each). A 4 mm plug of a filamentous fungus culture was placed on one side of the Petri dish and incubated until the mycelium of the control reached the opposite side of the plate, or up to 5 days in case of the slower-growing fungus *G. penicillata*. A white arrow indicates the location of the agar strip containing the stilbene or phenolic acid treatment. Putative beneficial fungi associated with *I. typographus* are shown over a blue background. The pathogen *T. harzianum* is shown over a dark red background for comparison, n = 6 plates. **b)** Fungal growth area relative to the DMSO control treatment. Different letters indicate significant differences among groups (Kruskal-Wallis rank-sum test followed by a Tukey post-hoc test, *p* < 0.05, n = 6 plates, n.s. = not significant).

Given that phenolic acids occur at comparatively higher levels in newly founded maternal tunnels rather than in older samples (Suppl. Fig. S6), we hypothesized that these yeast stilbene metabolites may also play a role in regulating fungal growth during the early stages of gallery colonization. We found that the biomass of bark beetle-associated fungi was significantly reduced when grown on PDA amended with concentrations of the phenolic acid mix in the range formed by *K. capsulata*, the yeast producing the highest levels of these substances. When tested at this higher concentration, these stilbene metabolites significantly reduced the biomass of both the pathogen *T. harzianum* and the bark beetle filamentous symbionts (Suppl. Fig. S8). Phenolic acids seem to have antifungal effects that may also influence the growth of bark beetle symbiotic fungi, especially during the earliest stages of gallery colonization when these compounds are more abundant (Suppl. Fig. S6). Curiously, the yeast *K. capsulata* possesses a specialized stilbene metabolism producing much higher levels of phenolic acids than the other species tested (SI Appendix – *Kuraishia capsulata* metabolism, Suppl. Figs. S9-S11). Thus, the occurrence of high amounts of phenolic acids in bark beetle galleries may depend on the relative abundance of *K. capsulata* in the yeast community.

## Discussion

The associations between scolytines and yeasts are widespread across geographic locations and beetle species (12,13,19,21). Yeast communities seem to be stable in certain bark beetle species, and over the years it has been speculated that these microorganisms perform beneficial functions for their insect partners (12,14). We provide first evidence that symbiotic yeasts may benefit bark beetles in a system that includes the host tree, the beetle’s filamentous symbionts, and fungal antagonists. Our results show that yeasts modify spruce defense chemicals, in a way that could protect the beetle’s eggs from a pathogen and regulate the growth of filamentous ectosymbionts in the galleries (Fig. 8). Furthermore, behavioral assays revealed that adult beetles are attracted to their symbiotic yeasts, and phylogenetic analysis showed these isolates are related to those obtained from other insects. However, the question of whether these associations are mutualistic remains unanswered. We speculate that beetle-mediated dispersal and early access to freshly excavated galleries could be beneficial for the yeasts, but whether these advantages go beyond phoresy remains uncertain.

**Fig. 8.**
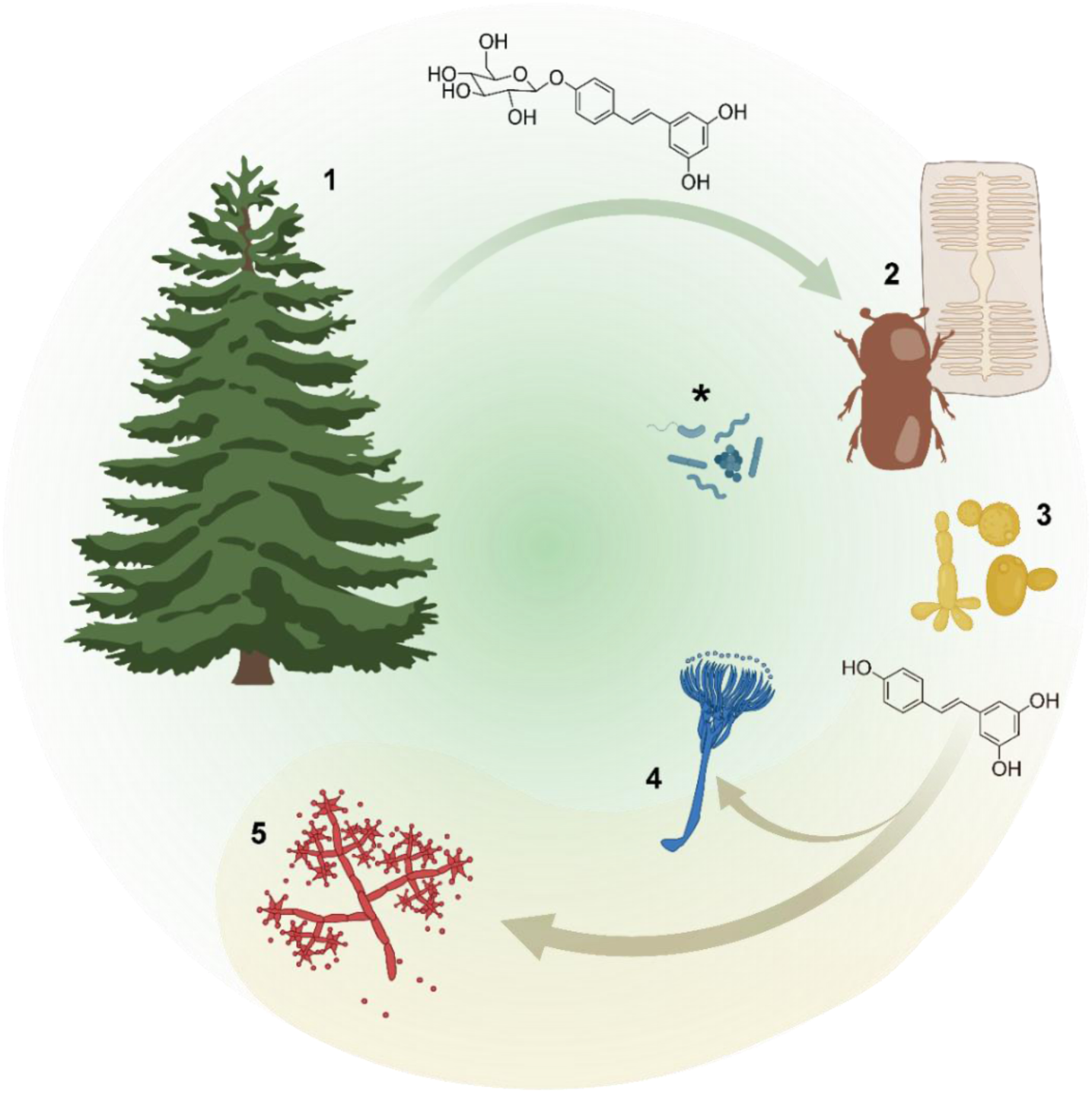
Schematic summary of the influence of yeasts on bark beetle-host tree interactions. (**1**) Norway spruce (*P. abies*) produces a diverse array of anti-herbivore and antimicrobial defense metabolites, including high levels of terpenoids and phenolic compounds. (**2**) Bark beetles (*I. typographus*) have evolved to colonize and feed on spruce despite its ample chemical defenses, occupying a niche where competition with other herbivores is low. One of the adaptations to such a hostile environment is symbiosis with microorganisms. Bark beetles associate with a suite of bacteria (*), yeasts and filamentous fungi, some of which are vectored by the insects into spruce during colonization. (**3**) Yeasts are among the first microorganisms in the ecological succession that occurs within the beetles’ galleries. *I. typographus*-associated yeasts have coopted one of spruce’s main defense chemical classes (stilbenes) and utilize them to mediate the growth of the insect’s filamentous fungal partners and inhibit the growth of non-beneficial fungi. Their niche is restricted to the galleries and the beetles themselves. (**4**) Filamentous fungi that are partners of the beetles typically increase in abundance in the galleries at later stages and colonize tree tissue beyond the limits of the galleries and the phloem. (**5**) Fungal competitors are challenged by the specialized adaptations of bark beetle symbionts to the trees’ chemistry, and are only late colonizers of the galleries. Once this occurs, the beetles are likely to have completed their life cycle and migrated to a new host, carrying their microbes with them. (**\***) While bacteria were not part of this study, their roles in the multi-trophic interactions that occur in the galleries are also likely to be relevant for the beetles and their filamentous fungal symbionts (15,18,73).

Similar to burying beetles, where vertically transmitted *Yarrowia* yeasts play an important role in preserving the insect’s feed and breeding substrate (32,33), bark beetles shape the microbial communities in the galleries to increase substrate quality for their offspring (18). In both systems, the beetles exhibit a subsocial parental care that involves substrate manipulation, vertical transmission of microbes, and some degree of grooming (34) (in *I. typographus*, grooming is limited to the maternal galleries and the phloem plugs). For both beetles, the preparation and preservation of their nutritional resources relies on a complex microbial community composed of fungi and bacteria that defend their feeding and breeding substrates against antagonists (33).

In the context of defensive symbioses, the strategies used by microorganisms to protect their animal hosts can be classified in four categories: host vigor improvement, competitive exclusion, immune stimulation or priming, and direct chemical defense (35). In insects, there are some well-known examples of symbiont-mediated chemical defense. This is the case for beewolf wasps (*Philanthus triangulum*) that cultivate *Streptomyces philanthi* in specialized antennal bacteriangia (36) and apply them to their brood cells before oviposition (37). The bacteria produce an antimicrobial mixture of streptochlorin and several piericidin derivatives, which offer the beewolves’ cocoons a prophylactic protection against opportunistic soil-dwelling pathogens (38). Another bacterium, *Burkholderia gladioli*, is vertically transmitted by female darkling beetles (*Lagria villosa*) to their eggs. Depending on the strain, the microbe protects its host’s offspring by deploying the antifungal lagriamide or a mix of antimicrobials including caryoynencin, sinapigladioside, lagriene, and other bioactive compounds (26,39). In both the beewolf and the *Lagria* symbiosis, the microorganisms synthesize various compounds that shield vulnerable life stages of their hosts from antagonists present in their breeding substrates.

As in the aforementioned examples, the bark beetle-associated yeasts *K. capsulata, K. molischiana* and *N. holstii* deploy ‘chemical cocktails’, mixtures of antifungal compounds, as a potential defensive strategy. The particularity of these yeasts, however, is that they do not produce these defensive substances *de novo,* but instead co-opt and modify Norway spruce defense compounds in a way that could protect the offspring of their insect host against pathogen attacks. This strategy is enabled by the host tree, which accumulates very high concentrations of antimicrobial compounds in the phloem tissues of its main trunk (40). Hence, when yeasts hydrolyze the spruce stilbene glucosides to their corresponding and more toxic aglucones, this leads to high concentrations of antimicrobial substances in bark beetle galleries. This metabolism is apparently not harmful to the beetles themselves, which have been reported to convert stilbene glucosides to aglucones via their own enzymes (27). However, this outcome has a profound negative impact on the tree by benefiting one of its main herbivores. Thus, a plant’s production of high levels of antimicrobial defenses may not necessarily be to its benefit if these are subsequently deployed against a pathogen of its herbivore. Critical for this outcome is also that the herbivore’s microbial symbiont, the yeast, must be resistant to the toxic effects of stilbene aglucones.

The ability of bark beetles to also transform stilbene glucosides to aglucones in their guts (27) may contribute to the antifungal activity in their galleries, especially at later stages of infestation when their galleries are filled with frass. However, the presence of yeasts in the early phases of gallery development suggests an important role for these symbionts in provisioning antimicrobial stilbenes during vulnerable stages of beetle development, although we cannot exclude the possibility that there are additional compounds involved in these protective effects, such as the terpenoid components of the resin, some of which are known to have antifungal properties (41). Nevertheless, our findings provide evidence for a novel mechanism of chemical defense in herbivore-microbe symbioses, where associates of an insect’s fungal community metabolize host plant defense compounds in a manner that may reduce pathogen pressure on the insect.

## Materials and Methods

### Culture media

Throughout our study, we used different culture media for yeast and filamentous fungi cultivation. For initial characterization (yeast isolation, tree defense metabolite quantification after yeast growth) we used spruce phloem agar medium (SPA) which consisted of 2% w/v freeze-dried *P. abies* phloem powder with 2% w/v bacteriological agar (Carl Roth®). For initial fungal confrontation assays, behavioral assays, and stilbene metabolism tests we used standard potato dextrose agar medium (PDA, Carl Roth®). For the rest of the confrontation assays we used enriched spruce phloem agar medium (rich SPA) which consisted 2% w/v freeze-dried *P. abies* phloem powder with 1% w/v bacteriological agar (Carl Roth®) amended with 1% w/v PDA to promote fungal growth. Liquid cultures for glycerol stocks and inoculum for the experiments were grown in potato dextrose broth (PDB, Carl Roth®).

### Yeast isolation and identification

We used a cultivation-based approach to characterize the yeast communities associated with the European spruce bark beetle. Adult *I. typographus* were collected in the summer of 2024 from pheromone-baited traps in the Tharandt forest (Saxony, Germany), where traps were emptied twice a day. Subsequently, live beetles were stored individually at 4°C in sterilized 1.5 mL tubes for max. 2 days, until they were further processed. In total, fifteen beetles were used for the isolation of yeasts. Each individual was placed in a microcentrifuge tube containing 500 µl of sterile phosphate buffer saline (PBS) and 0.1% Tween solution. The tube was vortexed for 20 seconds to obtain a suspension of the microorganisms present on the insect’s surface. Each beetle was subsequently removed from the tube and the head was excised from the body with a sterile scalpel. Each beetle segment was transferred to a microcentrifuge tube containing 500 µl of a solution of sterile PBS and 0.1% Tween 20®, macerated using sterile micro-pestles and vortexed vigorously for 20-40 seconds. A 100 µl aliquot of each sample was inoculated into spruce phloem agar (SPA) and incubated for 72 hours at 25°C and 65% relative humidity. Morphologically distinct colonies were picked from each plate, streaked onto potato dextrose agar (PDA, Carl Roth®) and incubated for 48 hours at 25°C and 65% relative humidity. This step was repeated until pure cultures were obtained. Glycerol stocks from all isolates were prepared and stored at −80°C for long-term preservation.

DNA was extracted from pure cultures using the Jena Bioscience Animal and Fungi DNA preparation kit following the manufacturer’s instructions. DNA quality and yield were measured using a nanospectrophotometer (Nanodrop 2000c, Thermo Fischer Scientific). The large subunit gene (LSU, 28S rRNA) was amplified with a Polymerase Chain Reaction (PCR) using the primers LROR (GTACCCGCTGAACTTAAGC) and LR5 (ATCCTGAGGGAAACTTC) (42). The PCR program consisted of an initial denaturation step of 4 minutes at 95°C followed by 30 cycles of 45 s at 94°C, 30 s at 55 °C and 60 s at 72°C, and a final extension step at 72°C for 6 minutes. The resulting products were purified using the Wizard SV Gel and PCR clean-up system (Promega). The purified amplicons were sequenced bidirectionally in a 3730XL DNA Analyser (Applied Biosystems from Thermo Fisher Scientific). The sequences were analyzed using SnapGene Viewer v6.1. The reverse sequences were transformed to their reverse-complementary strands and aligned with the forward sequences to create the consensus sequences using GeneDoc v2.7. These final sequences were compared against the NCBI database (43) and the taxonomy was assigned to the species level (>99% identity match). A representative isolate of each species was used for the subsequent experiments.

### Yeast phylogeny

To infer the phylogenetic affiliation of the yeast taxa obtained in the study, we performed maximum likelihood analyses using R (version 4.5.1) (44) and IQ-Tree (version 2.2.2.) (45). The phylogeny comprises 37 yeast species using the large subunit ribosomal ribonucleic acid (LSU; 28S rRNA), from which nine sequences were generated in this study while the remaining sequences were retrieved from the NCBI GenBank database (Suppl. Table S1).

We systematically compared multiple sequence alignment methods to identify the optimal alignment for downstream phylogenetic analyses using the package “msa” (46). Four algorithms were tested: Clustal Omega(47), ClustalW (48), Muscle (49), and MAFFT (50). To improve alignment quality by removing poorly aligned or ambiguously aligned regions, Gblocks Version 0.91b(51) was applied to each alignment filtering out positions with high gaps or low conservation. Next, each alignment and its corresponding Gblocks-filtered version were assessed using four quality metrics: mean Shannon entropy per position, gap percentage, proportion of fully informative positions, and log-likelihood values derived from fitting phylogenetic trees using maximum likelihood with neighbor-joining starting trees applying the packages “Biostrings”, “entropy”, “seqinr”, and “phangorn” (49,52–58). The best Gblocks-filtered version (in this case Muscle) was selected based on the highest overall quality scores. Then, we used the model finder implemented in IQ-Tree to select the best-fitting model (TIM3+F+I+R3) for further analyses. Phylogenetic trees were inferred using maximum likelihood with robust branch support estimation, including 8,000 ultrafast bootstrap replicates and Shimodaira-Hasegawa approximate likelihood ratio test (SH-aLRT) branch tests, providing support values for the inferred topology and clades (59). We rooted calculated trees based on a fungal outgroup (*Sugiyamaella* spp.) using the “ape” package (60). The packages “dplyr”, “tidyverse”, and “stringr” were used for additional data processing and filtering (61–63); “ggtree”, FigTree and Adobe Illustrator (version 26.5) were used to visualize the final maximum likelihood tree (64,65).

### Behavioral assays

To examine whether adult *I. typographus* were attracted to the yeast isolates, we performed behavioral assays with a previously described method (66), where adult beetles confined to a large Petri dish choose among samples based only on olfactory cues. We used beetles collected from pheromone-baited traps in the Tharandt forest (Saxony, Germany) during 2024. Here, we included six *I. typographus*-associated yeasts, the yeast *D. ontarioensis*, a symbiont of another Norway spruce bark beetle (25), and *Trichoderma harzianum*, a pathogen in the bark beetle gallery environment (Suppl. Table S2).

To prepare the arenas, 100µl of liquid yeast culture (OD600 = 0.1) were inoculated on PDA for 48 hours at 25°C and 65% humidity. Then, a 6 mm Ø plug was taken from the inoculated PDA plate and used as one of the choice samples, and a non-inoculated PDA plug was used as an alternative choice. For the filamentous fungus *T. harzianum*, we used one plug (6mm Ø) from a 4-day old culture growing on PDA. We added four adult beetles to each and randomized all individual assay arenas to exclude site-specific effects. The arenas were transferred to a laminar hood in darkness at 25°C, where choices were monitored every second hour for up to six hours. To test each fungus, we used a total of 10 plates containing four beetles each (n=40).

### Egg infection bioassay

An existing egg infection bioassay for tenebrionid beetles(26) was adapted for the bark beetle system (Suppl. Fig. 5). In brief, the wells of a 96-well plate were filled to ⅓ of their volume with autoclaved vermiculite and sterile distilled water. Autoclaved filter paper discs were placed in each well, and 1.5 µl of a *T. harzianum* spore suspension (5-10 spores/µl) was inoculated on each disc. Autoclaved freeze-dried *P. abies* phloem was rehydrated with autoclaved water until a loose dough consistency was obtained. Phloem spheres (hereafter artificial plugs) of approximately Ø 3 mm were formed using a sterile spatula and one of them was placed on top of each filter disc with sterile forceps. *I. typographus* eggs were collected from the laboratory rearing in Jena, Germany, surface-sterilized following the protocol established by Peng *et al.*(67) and randomly assigned to sterile or *Kuraishia capsulata*-inoculated (10^4^ CFU/plug) artificial plugs. Non-sterilized eggs carrying their native microbiota were placed on sterile artificial plugs as a negative control. A total of 18 replicates were carried out per treatment. The plate was sealed with parafilm and incubated at 25°C and 65% relative humidity in the dark for five days. A blinded assessment of the infection status was carried out daily. A sample was marked as infected when mycelium of *T. harzianum* was visible on the artificial plug or the egg under a stereomicroscope (Leica, Germany).

### *In vitro* confrontation assays

Liquid yeast cultures (*N. holstii*, *W. bisporus*, *K. molischiana*, *K. capsulata*, *Y. scolyti*, *Y. tenuis;* OD600 = 0.1) were pipetted on PDA or rich SPA (spruce phloem agar enriched with 1% PDA) plates forming a straight line along the diameter of the Petri dish (Figure 4). A Ø 6 mm plug of filamentous fungal culture (*Trichoderma harzianum, Endoconidiophora polonica* or *Grosmannia penicillata*) was placed at one of the opposing ends of the plate. The fungi were co-cultivated for up to two weeks at 25°C and 65% relative humidity. Pictures were taken every day with an EOS 600D (Canon) camera to record fungal growth and the presence/absence of inhibition zones. Filamentous fungi inoculated in the absence of yeasts were used as a control. Six replicates were carried out per media-yeast-filamentous fungus combination.

### Stilbene quantification *in vitro* and in bark beetle galleries

For the *in vitro* analysis, 100 µl of liquid yeast cultures (*N. holstii*, *W. bisporus*, *K. molischiana*, *K. capsulata*, *Y. scolyti*, *Y. tenuis;* OD600 = 0.1) were inoculated on SPA plates supplemented with a sterile cellophane disk (NeoLab, Germany) to facilitate biomass collection and incubated for five days at 25°C and 65% relative humidity. Non-inoculated spruce phloem agar was sampled at the start of the experiment and after five days of incubation to be used as negative controls. The yeast biomass and the agar were sampled separately and freeze-dried or processed immediately after collection. Methanol extracts were prepared from dried yeast biomass (sample dry mass range 20-40mg) by homogenizing it with 1 mL of methanol (Honeywell) and three metal beads (Ø 3 mm, Askubal) on a paint shaker (Skandex SO-10M, Fluid Management Europe, The Netherlands) for 5min. Samples were centrifuged afterwards at 9400 rcf for 5 min and the supernatant was collected for further analyses. Stilbenes and phenolic acids were quantified with an LC-MS/MS system. For the gallery samples, nine 25 cm-long Norway spruce logs were placed in individual rearing cages and exposed to 40 adult beetles each. They were kept in a walk-in rearing chamber (Viessmann, Germany) with 16 h of light and 8 h of darkness at a temperature of 25°C and relative humidity of 60%. The different parts of the galleries -maternal gallery, phloem plugs, larval galleries, pupal chambers- were sampled from three logs per timepoint (5-, 14- and 21-days post-infestation). Four to six independent galleries were sampled in total per timepoint. In the case of the phloem plugs, all the plugs present along one maternal gallery were pooled to create a sample. A sample of the unattacked adjacent phloem (5 cm away from the beetle galleries) was included for each sampled gallery. The samples were freeze-dried immediately after each sampling batch. Methanol extracts were prepared following the protocol indicated earlier. We performed liquid chromatography-mass spectrometry (LC-MS/MS) on an Agilent 1200 HPLC system (Agilent Technologies, Boeblingen, Germany) coupled to an API 3200 tandem mass spectrometer (Applied Biosystems, Darmstadt, Germany). A detailed description of the methods is available in the SI Appendix - Methods. Samples were normalized by the individual sample dry weight unless fresh samples were extracted directly. In these cases, the fresh weight was used to normalize the concentrations.

### Break-down of phenolic acids by filamentous fungi and yeasts

To test the capabilities of filamentous fungi (n = 8, Suppl. Table 2) and yeasts (n = 7) to break down protocatechuic acid, vanillic acid, and 4-hydroxy benzoic acid, we supplemented PDA with an artificial blend containing each of the three phenolic acids originally dissolved in dimethyl sulfoxide (DMSO, Sigma) to a final concentration of 300 µg/mL of culture media (resulting final concentration of DMSO in the media was 2%). As a control served culture media supplemented with 2% DMSO. Additionally, we added a sterilized cellophane disk to separate biomass from culture medium. Yeasts were first subcultured overnight in liquid potato dextrose broth (PDB, Roth) at 25°C. Then, we inoculated 100 µl of the yeast suspension with an OD600 of 0.1 on each plate. Filamentous fungi were first sub-cultured on PDA and a plug of actively growing mycelium (6 mm Ø) was transferred on our experimental medium. All plates (n = 6 per isolate) were incubated at 25 °C and 65% humidity for up to 4 days for single-celled yeasts and for up to 12 days for filamentous fungi. Then, we removed the cellophane disk and took pure medium samples, which were subsequently freeze-dried and processed for further LC-MS/MS analyses using methanolic extracts as described in the previous section.

### Agar strip antifungal activity assays

Two-centimeter-wide strips were removed from the middle of PDA plates to create a well in the agar that crossed the full length of the diameter of each plate. The well was subsequently filled with PDA medium amended with one of the following treatments: DMSO (2 %v/v DMSO/PDA medium, control treatment), a stilbene aglycone mix dissolved in DMSO (piceatannol, resveratrol and isorhapontigenin at a final concentration 500 μg/ml PDA medium each), or a phenolic acid mix dissolved in DMSO (protocatechuic acid, vanillic acid and 4-hydroxybenzoic acid at a final concentration of 500 μg/ml PDA medium each) A 6 mm plug of filamentous fungus culture (*T. harzianum, E. polonica* or *G. penicillata;* n = 6) was placed on one side of the Petri dish and incubated until the mycelium of the control treatment reached the other end of the plate. The growth of each treatment was recorded with a Canon digital reflex camera attached to a still stand situated at 40 cm above the plates. Total growth area was measured with ImageJ (version 1.54p).

### Filamentous fungi growth in the presence of phenolic compound mixes

Five filamentous fungi (*O. bicolor, E. polonica, G. penicillata, C. minuta,* and the pathogen *T. harzianum*, n = 6) were grown in the presence of phenolic compound mixes. The treatments consisted of PDA medium amended with one of the following treatments: DMSO (2 %v/v DMSO/PDA medium, control treatment), a stilbene aglycone mix dissolved in DMSO (piceatannol, resveratrol, and isorhapontigenin at a final concentration 250 μg/ml PDA medium each), or a phenolic acid mix dissolved in DMSO (protocatechuic acid at a final concentration of 2000 μg/ml PDA medium; vanillic acid and 4-hydroxybenzoic acid at a final concentration of 800 μg/ml PDA medium each). The plates were supplemented with a sterile cellophane disk (NeoLab, Germany) to facilitate biomass collection, and Ø 6mm plugs of mycelium were used as the inoculum. To account for the different growth rates of each fungal species, the biomass was harvested at the time point when the mycelium of the control treatment had covered the plate, except for *C. minuta*, where it was collected after 14 days. The mycelium was collected with a cell scraper, freeze-dried and weighted.

### Resveratrol metabolism in *Kuraishia capsulata*

To investigate the resveratrol metabolism in *K. capsulata*, 100 µl of liquid yeast culture (*W. bisporus*, *K. molischiana*, OD600 = 0.1) were inoculated on PDA amended with either DMSO (2 %v/v DMSO/PDA medium, control treatment) or resveratrol dissolved in DMSO to a final concentration of 200ug resveratrol/g PDA medium (n = 6 per yeast per treatment), all supplemented with a sterile cellophane disk (NeoLab, Germany) to facilitate sampling the agar without yeast biomass. The plates were incubated for 72 hours at 25°C and 65% relative humidity. A sample of agar was taken from each plate, freeze dried and extracted in methanol following the protocol described earlier.

Ultra-high-performance liquid chromatography–electrospray ionization– high resolution mass spectrometry (UHPLC–ESI–HRMS) was performed on the methanol extracts with a Dionex Ultimate 3000 series UHPLC (Thermo Scientific) and a Bruker timsToF mass spectrometer (Bruker Daltonik, Bremen, Germany). The detailed methodology is available in Suppl. File 4. After identification of 3,5-dihydroxybenzoic acid as a unique breakdown product of resveratrol in the agar inoculated with *K. capsulata,* we repeated the cultivation experiment using this phenolic acid as a supplement to the growth media (Sigma Aldrich, Germany). In brief, 100 µl of liquid yeast culture (*W. bisporus*, *N. holstii, K. molischiana*, OD600 = 0.1) were inoculated on PDA amended with 3,5-dihydroxybenzoic acid dissolved in DMSO at a final concentration of 50 μg/ml PDA medium (n = 5 per yeast) supplemented with a sterile cellophane disk (NeoLab, Germany). The plates were incubated for 5 days at 25°C and 65% relative humidity. A sample of agar was taken from each plate, freeze dried and extracted in methanol as described earlier. The extracts were used to perform untargeted metabolite analysis as described above.

### Statistical analyses

All statistical analyses were carried out using R version 4.5.1.(44) The behavioral choice assays were analyzed using binomial tests on pooled data. The number of insects choosing each fungus was summed across all trials, and the resulting proportion of individuals selecting the fungus was compared to a null expectation of 0.5. The infection probability in the egg bioassays was estimated using a Kaplan-Meier estimate followed by a log-rank test and a Cox proportional hazards regression with the packages “survival” and “ggsurvfit”.(68,69) Significant differences among treatments in metabolite concentrations, fungal growth area and biomass were assessed with a Kruskal–Wallis rank-sum test using “biostat”.(70) Dunn’s test was used to calculate pairwise comparisons between the sample types using Bonferroni’s correction for multiple sample comparison. Heatmaps were created with the package “pheatmaps”.(71) Plots were created using the package “ggplot2”(72) and edited for improved visualization in Adobe Illustrator version 26.5. Molecular structures were drawn using ChemDraw version 23.1.2.7. Illustrations were created with Biorender.com.

## Author Contributions

A.B.Q., M.K., J.G. and M.L. designed the experiments. L.S.P. and M.L. performed yeast isolation, molecular characterization, and initial stilbene profile analysis. A.B.Q. performed egg surface sterilization and in vivo antifungal bioassays, fungal growth inhibition assays and confrontation assays. M.L. and A.B.Q. performed stilbene degradation assays. M.L. carried out behavioral olfaction-guided bioassays and phylogenetic analysis. M.R. carried out untargeted chemical analysis. R.S. provided astringin for the standards. Experiments and data analyses were carried out with input from M.R., R.S., M.K., J.G, and ML. J.G., M. L. and M.K. provided supervision and acquired funding. A.B.Q., L.S.P., M.R. and M.L. wrote the manuscript. All authors commented on the final draft of the manuscript.

## Data availability

The sequences of the isolates obtained in this study are available in NCBI – accession numbers PZ239427 to PZ239550.

## Acknowledgments

We would like to thank Natascha Rauch for her valuable assistance in the laboratory and the Forstamt Jena-Holzland for facilitating tree felling for our experiments. We are grateful for the financial support from the Max Planck Society, the European Research Council through an ERC Consolidator Grant to MK (ERC CoG 819585 “SYMBeetle”), and the German Research Foundation to ML (DFG: Grant number 520486374).

