## Supplemental Information for "Symbiotic yeasts of a bark beetle transform major tree defenses into beetle protectants"

**This PDF file includes:**

Figures S1-S14
Tables S1-S2
Supporting text: *Kuraishia capsulata* metabolism
Supporting text: Methods

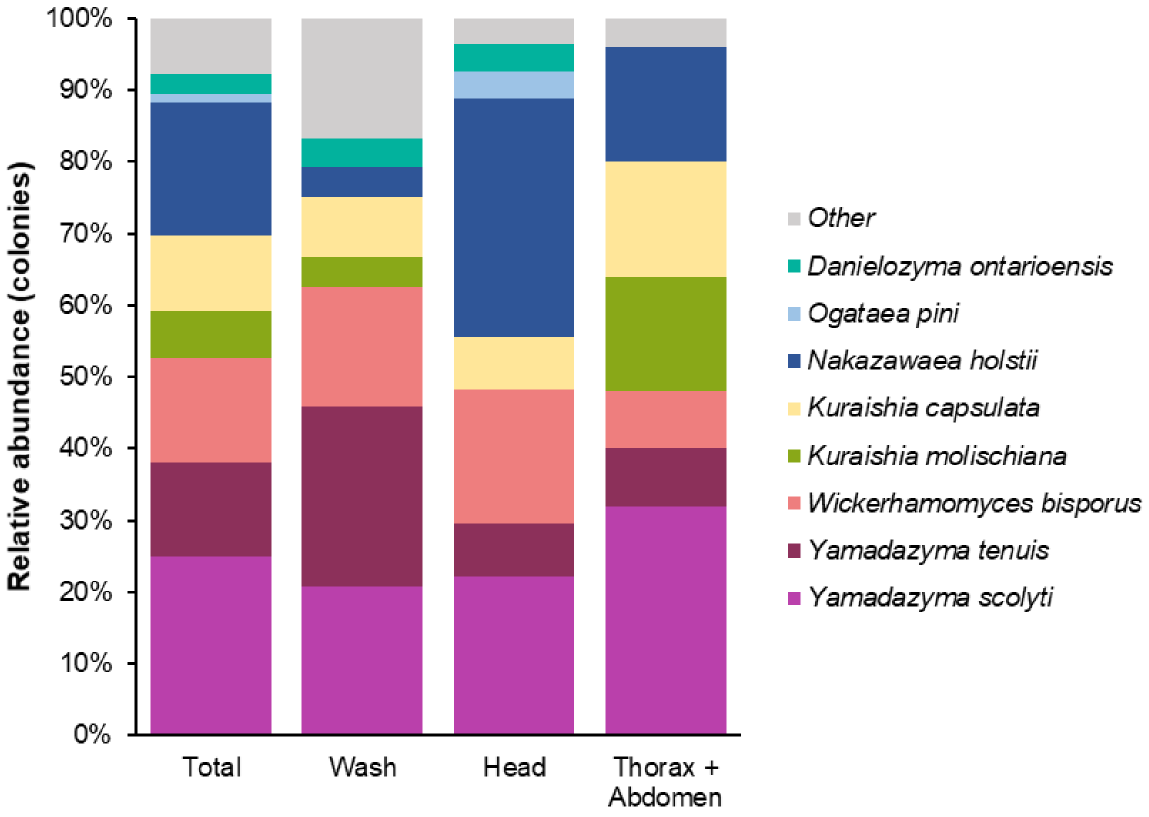


**Suppl. Fig. S1 | Relative abundances of yeasts isolated from adult *I. typographus*.** Yeast species isolated from the body surface (wash), heads, and abdomens of wild-caught *I. typographus* adults. The number of isolates of each taxon obtained from the beetles was used to calculate the relative abundance of each yeast species, n = 15 beetles.

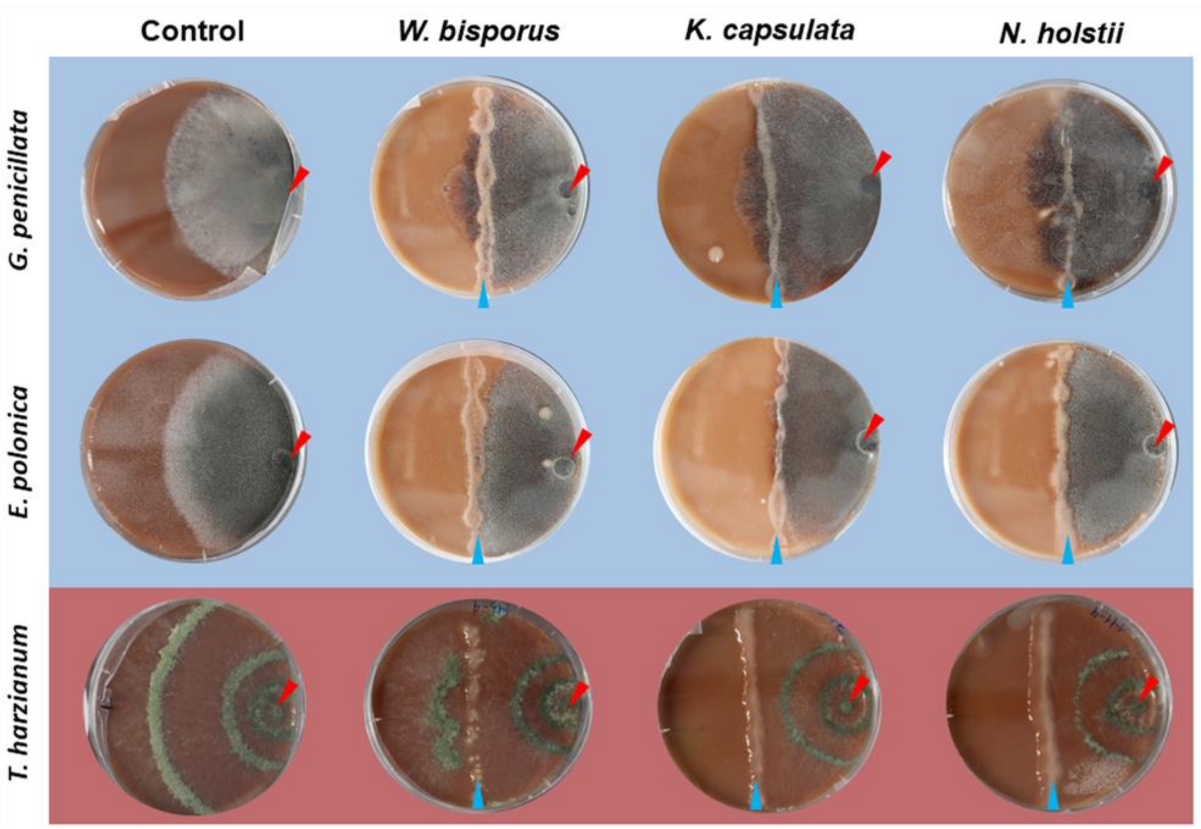


**Suppl. Fig. S2 | Confrontation assays of selected yeasts against bark beetle-associated fungi.** A yeast suspension (names listed across the top) was pipetted on an agar plate forming a straight line crossing the center. A 4mm plug of a filamentous fungus culture (names listed on the left side) was placed at one of the opposing ends of the plate. They were co-cultivated for five days on rich spruce phloem agar medium (2% SPA amended with 1% PDA). A blue arrow indicates the inoculation site of the yeast and a red arrow the placement of the filamentous fungi. Fungi associated with *I. typographus* are shown over a blue background, the pathogen *T. harzianum* is shown with a dark red background for comparison. Filamentous fungi grown in the absence of yeasts were used as a control, n = 6 plates.

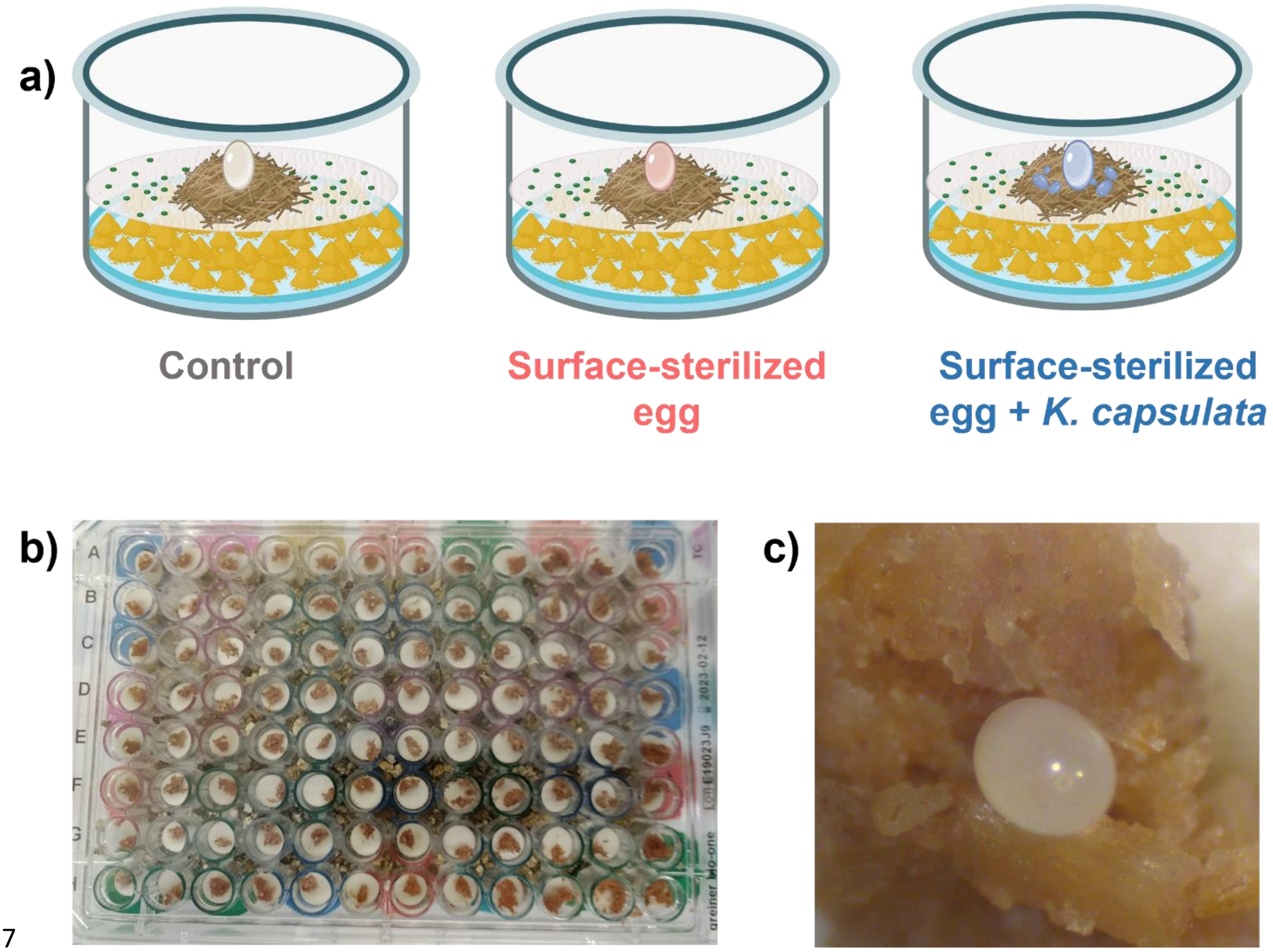


**Suppl. Fig. S3 | Egg infection bioassay setup. a)** Schematic view of the assay. The wells of a 96-well plate were filled to ⅓ of their volume with autoclaved vermiculite (depicted in yellow) and sterile distilled water. Autoclaved filter paper discs were placed in each well, and a *T. harzianum* spore suspension was inoculated on each disc. An artificial phloem plug (brown) was placed on top of each filter disc. Surface-sterilized eggs were randomly assigned to sterile or *Kuraishia capsulata*-inoculated artificial phloem plugs. Non-sterilized eggs carrying their native microbiota were placed on sterile artificial plugs as a control. **b)** 96-well plate with the eggs and phloem plugs. **c)** Detail of an egg on an artificial phloem plug inside of a well in a 96-well plate.

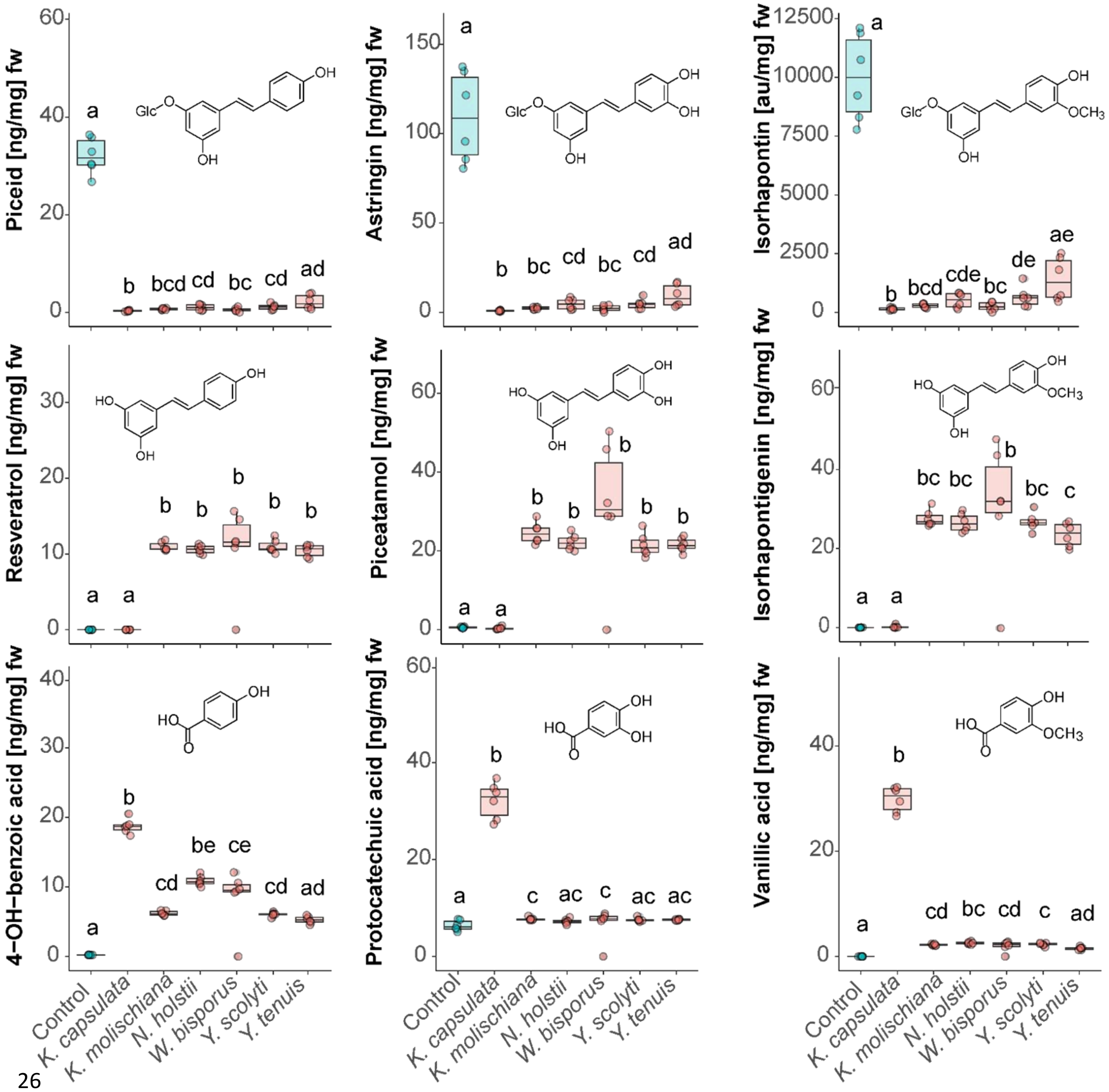


**Suppl. Fig. S4 | Stilbene and phenolic acid content in fresh spruce phloem agar after yeast growth.** The concentration of stilbene glycosides (top row), their aglycones (middle row) and the corresponding phenolic acids (bottom row) were measured in fresh spruce phloem agar medium five days after inoculation with *I. typographus*-associated yeasts. Non-inoculated spruce phloem agar was incubated during five days and used as a negative control. Concentrations are shown in nanograms per milligram of fresh weight except for isorhapontin, where the peak area was used instead (area units per milligram of fresh weight). Different letters indicate significant differences among groups (Kruskal-Wallis followed by Dunn's post-hoc test, $p < 0.05$, n = 6). In isorhapontin: "au/mg" = peak area/mg.

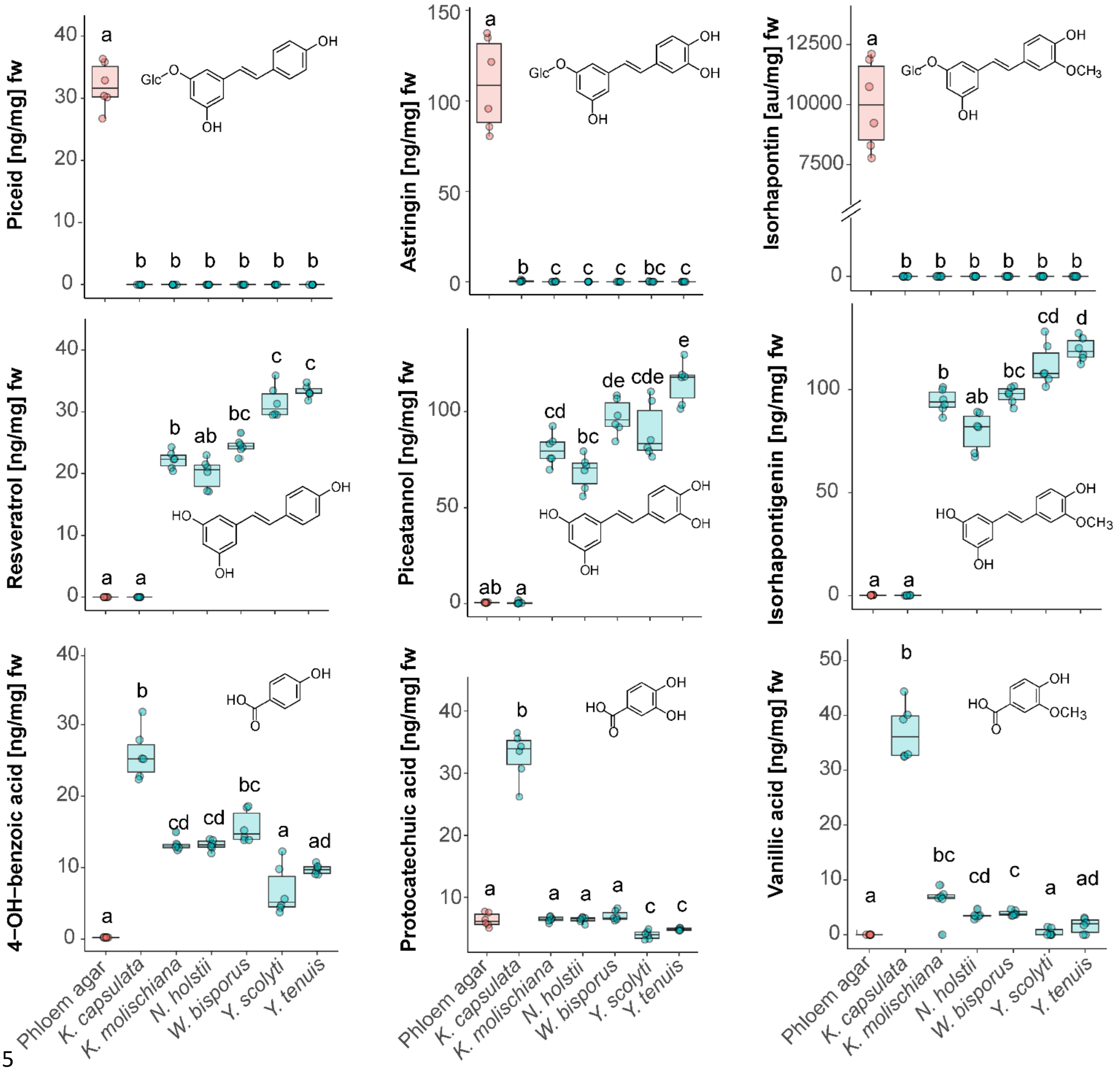


**Suppl. Fig. S5 | Stilbene and phenolic acid content in the fresh biomass of yeasts growing on phloem agar.** The concentration of stilbene glycosides (top row), their aglycones (middle row) and the corresponding phenolic acids (bottom row) were measured in the fresh biomass of yeasts that were grown on spruce phloem agar for five days. Non-inoculated spruce phloem agar was incubated during five days and used as a negative control. Concentrations are shown in nanograms per milligram of fresh weight except for isorhapontin, where the peak area was used instead (area units per milligram of fresh weight). Different letters indicate significant differences among groups (Kruskal-Wallis followed by Dunn's post-hoc test, $p < 0.05$, n = 6). In isorhapontin: "au/mg" = peak area/mg.

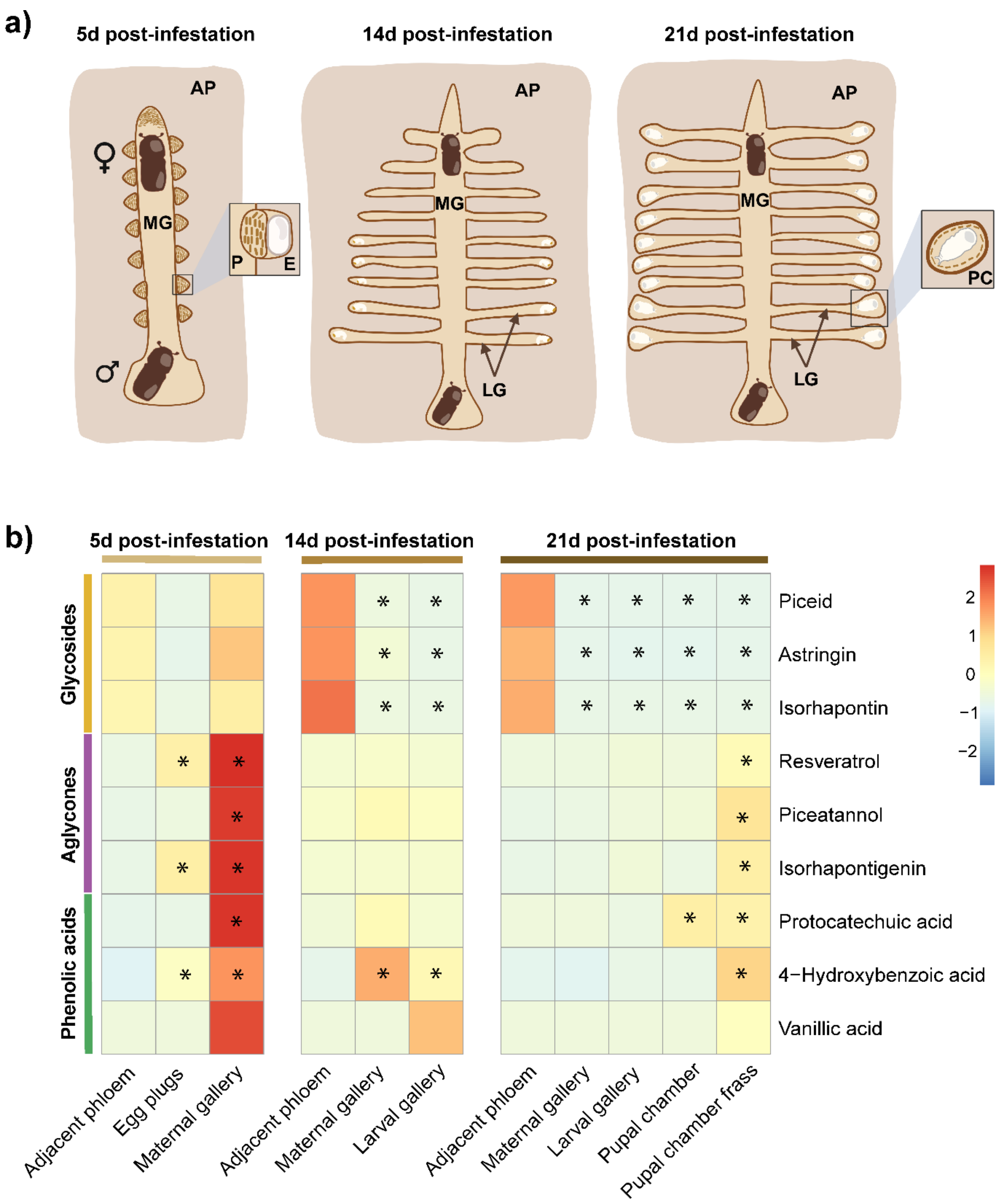


**Suppl. Fig. S6 | Stilbene and phenolic acid content in the galleries at different developmental timepoints. a)** Schematic view of gallery development 5, 14 and 21 days after infesting spruce logs with adult bark beetles. MG = maternal gallery, AP = adjacent phloem, P = plug, E = egg, LG = larval gallery, PC = pupal chamber. **b)** Heatmap of the average concentrations of stilbene glycosides, stilbene aglycones and phenolic acids present in different parts of the galleries at the sampled time points. The non-damaged phloem adjacent to the bark beetle galleries was included as a control on each sampling time point. Asterisks (*) indicate fields that were significantly different from their respective adjacent phloem control (Kruskal-Wallis rank-sum test followed by a Tukey post-hoc test, $p < 0.05$, n = 4-6). Isorhapontin concentration is reported in peak area/mg, while the rest of the compounds are ng/mg.

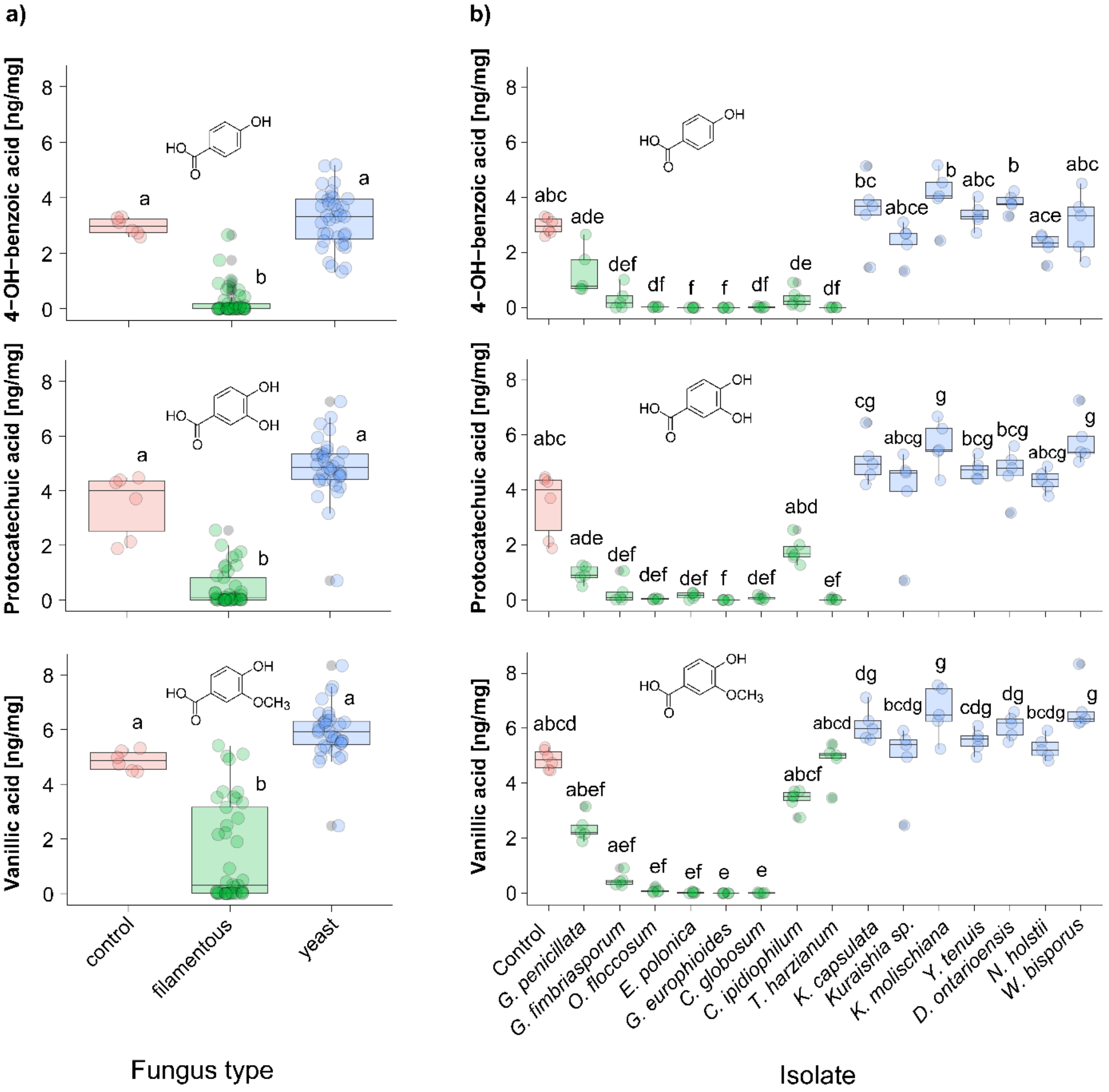


**Suppl. Fig. S7 | Phenolic acid metabolism by filamentous fungi and yeasts associated to bark beetles.** 4-hydroxybenzoic acid, protocatechuic acid and vanillic acid content in PDA supplemented with a mix of the three phenolic acids after incubation with filamentous fungi and yeasts associated to bark beetles. **a)** Aggregated data per fungus type. **b)** Phenolic acid content in the medium per isolate. Different letters indicate significant differences among treatments (Kruskal-Wallis followed by a Dunn's post-hoc test, $p < 0.05$, n = 6).

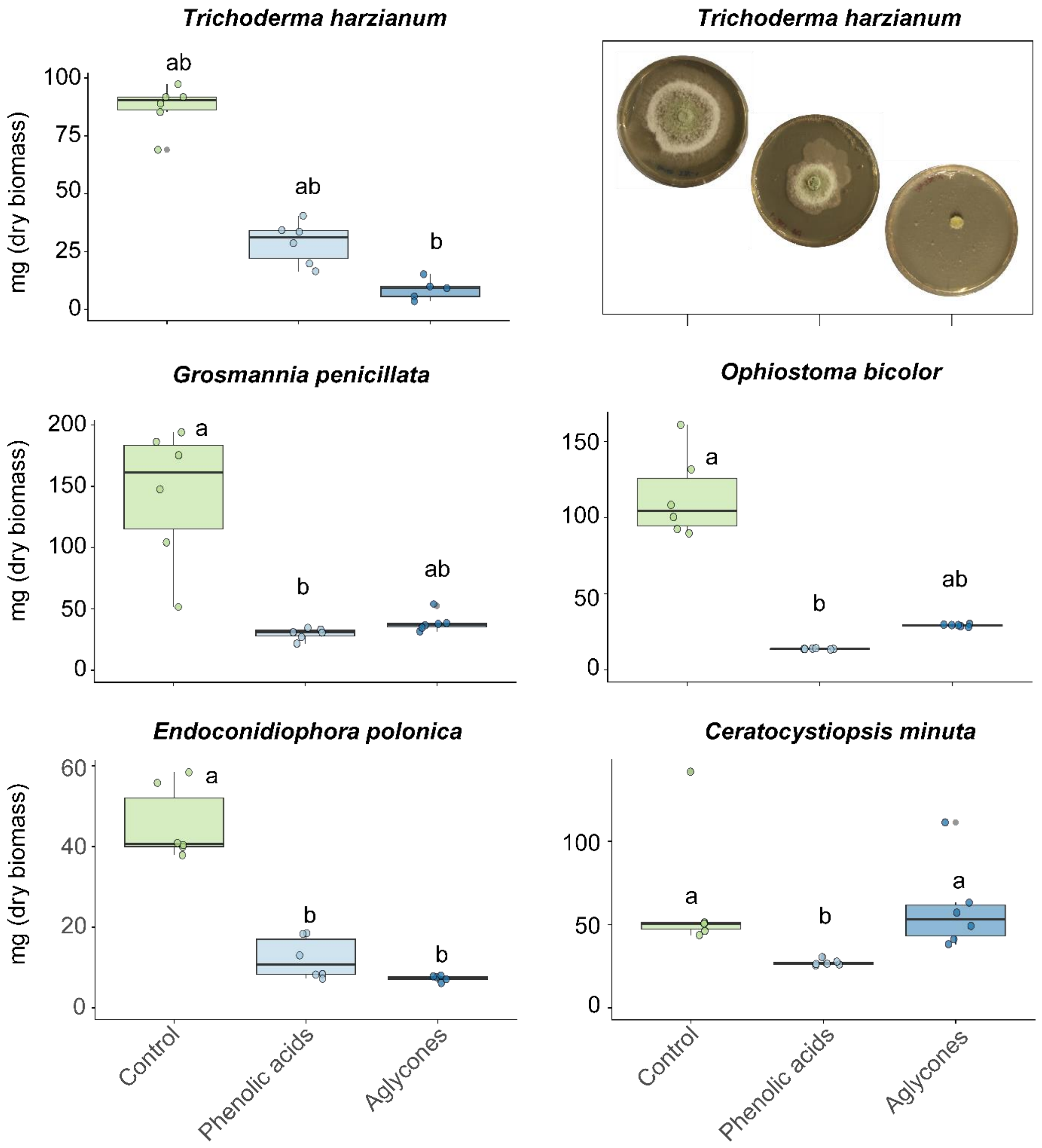


**Suppl. Fig. S8 | Fungal biomass grown on PDA supplemented with stilbene compound mixes or phenolic acid mixes.** Dry biomass of the saprophyte *T. harzianum* and four *I. typographus* filamentous symbionts grown on PDA amended with DMSO (2% DMSO, control treatment), a stilbene aglycone mix (piceatannol, resveratrol and isorhapontigenin at 250µg/g medium each), or a phenolic acid mix (protocatechuic acid 2000µg/g; vanillic acid and 4-hydroxybenzoic acid at 800µg/g medium each). To correct for the different growth rates for each species, the fungi were collected at the time point when the mycelium of the control treatments had covered the plate, except for *C. minuta*, where the biomass was collected after 14 days. Different letters indicate significant differences among treatments (Kruskal-Wallis followed by a Dunn's post-hoc test, $p < 0.05$, n = 6). Top right insert: representative pictures of *T. harzianum* growth on each medium 48 hours post inoculation.

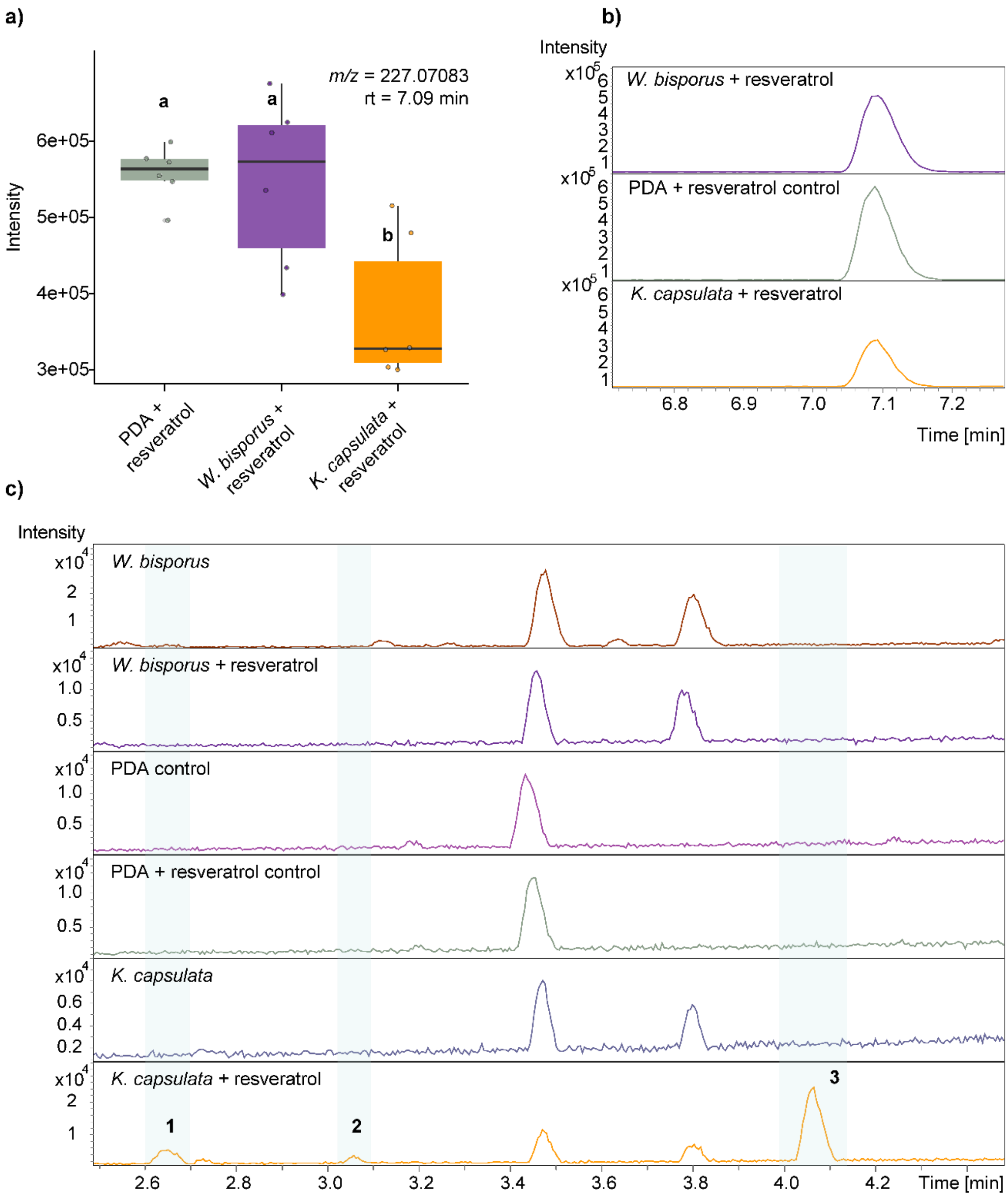


**Suppl. Fig. S9 | *K. capsulata* produces phenolic acids from resveratrol. a)** Resveratrol content (based on intensity of m/z 227.07083 from LC-Q-TOF-MS in negative mode) in resveratrol-supplemented PDA medium after 72 hours of *W. bisporus* and *K. capsulata* growth. Non-inoculated resveratrol-supplemented PDA incubated for 72 hours was used as a control. Different letters indicate significant differences among the treatments (Kruskal-Wallis followed by a Dunn's post-hoc test, $p < 0.05$, n = 6). **b)** LC-QTOF-MS representative total ion chromatograms in negative mode showing resveratrol peaks (*m/z* = 227.07083) in resveratrol-supplemented PDA after 72 hours of *W. bisporus* and *K. capsulata* growth. **c)** LC-QTOF-MS representative total ion chromatograms showing compounds identified in resveratrol-supplemented PDA after 72 hours of *K. capsulata* growth: (**1**) hydroxylated form of 3,5-dihydroxybenzoic acid, *m/z* = 169.01435; (**2**) 3,5-dihydroxybenzoic acid, *m/z* = 153.01943; (**3**) 4-hydroxybenzoic acid, *m/z* = 137.0244.

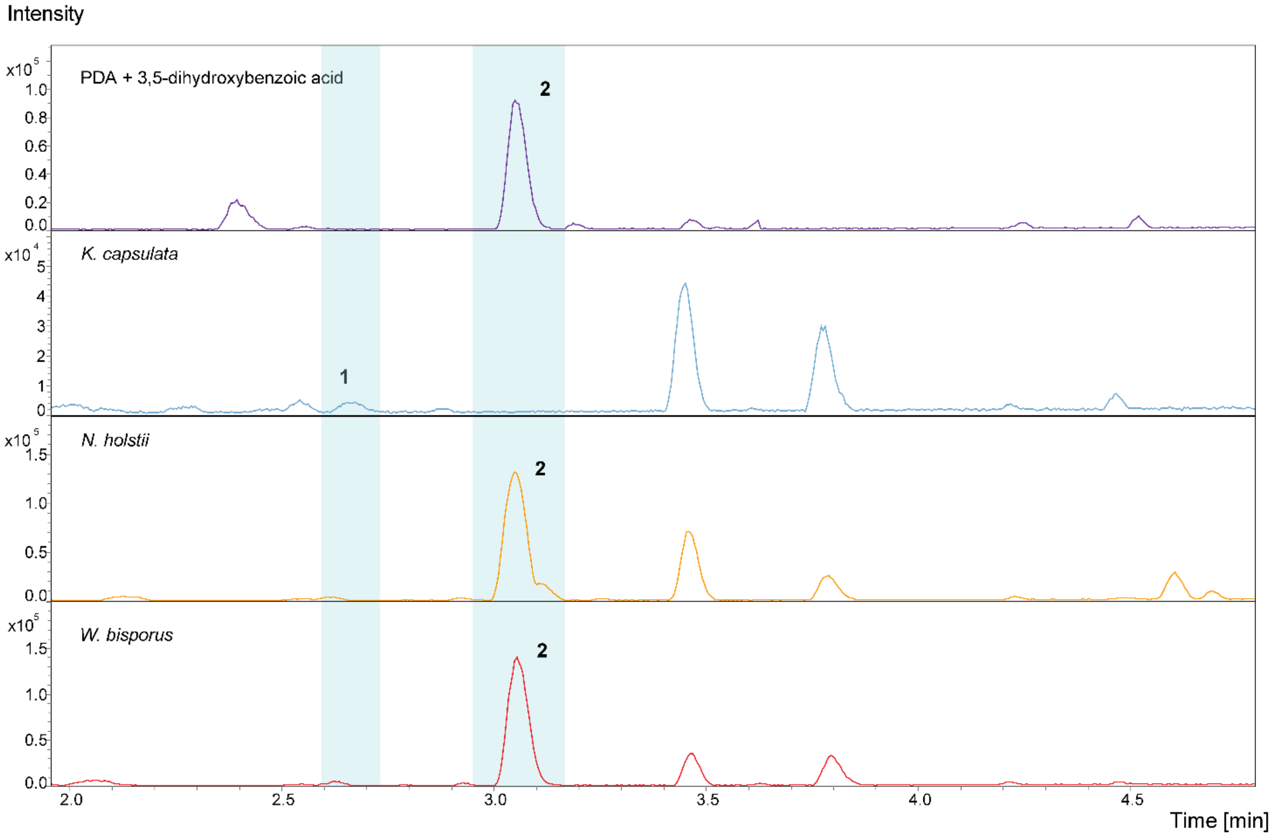


**Suppl. Fig. S10 | *K. capsulata* metabolizes 3,5-dihydrobenzoic acid.** LC-QTOF-MS representative total ion chromatogram showing compounds identified in 3,5-dihydroxybenzoic acid-supplemented PDA medium after a 72-hour incubation with *K. capsulata*, *W. bisporus* or *N. holstii*: (**1**) hydroxylated form of 3,5-dihydroxybenzoic acid, *m/z* = 169.01435; (**2**) 3,5-dihydroxybenzoic acid, *m/z* = 153.01943. Non-inoculated 3,5-dihydroxybenzoic acid-supplemented PDA incubated for 72 hours was used as a control (n = 5).

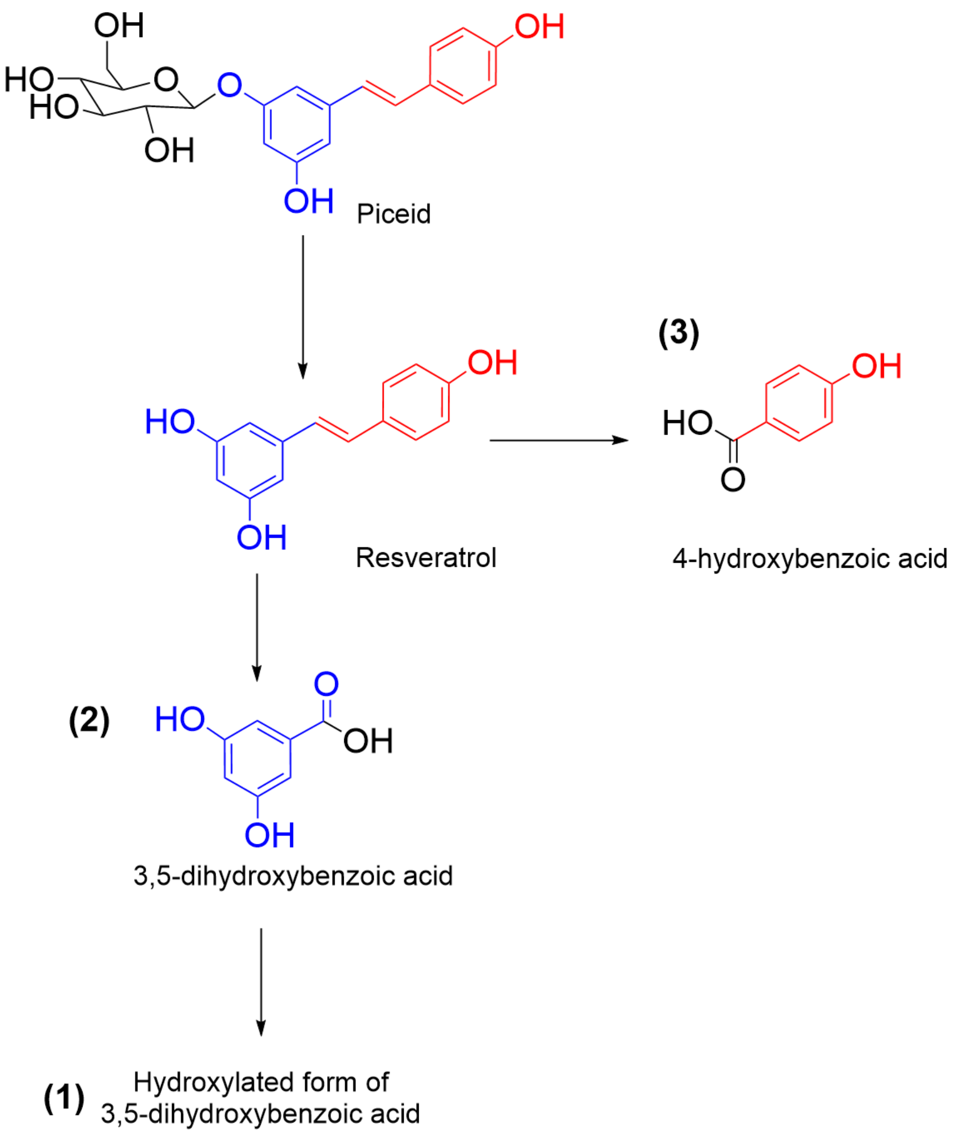


**Suppl. Fig. S11 | Proposed resveratrol metabolism in *K. capsulata*.** Piceid is deglycosylated into resveratrol. An oxidation step degrades the molecule into 4-hydroxybenzoic acid and 3,5-dihydroxybenzoic acid, which is further hydroxylated. The compounds identified in the LC-QTOF-MS analysis are indicated with the corresponding numbers: (**1**) hydroxylated form of 3,5-dihydroxybenzoic acid, (**2**) 3,5-dihydroxybenzoic acid, (**3**) 4-hydroxybenzoic acid.

**Suppl. Table S1:** Overview of all 37 yeast species used for phylogenetic analysis with the specimen identification number (ID) and the corresponding GenBank accession numbers for the LSU marker. (*****)Species from which we sequenced the LSU-region and which we used for phylogeny reconstruction.

| **Yeast species** | **Accession_N**o. | **Strain_ID** | **Isolate origin** | **origin category** |
|---|---|---|---|---|
| *Candida albicans* | NG054826.1 | | Skin lesion of *erosio interdigitalis* | human |
| *Candida orthopsilosis* | NG054816.1 | | Central venous pressure catheter tip inserted in human | human |
| *Candida tropicalis* | PP972331.1 | | Cauvery River water | water |
| *Cyberlindnera sp.** | | Y23 | *Ips typographus*, this study | *Ips typographus* |
| *Cyberlindnera xylebori* | MG674003.1 | | Tunnel of beetle *Xyleborus* sp. in *Fagus crenata* | insect |
| *Danielozyma ontarioensis** | | Y139 | *Ips typographus*, this study | *Ips typographus* |
| *Danielozyma sp.* | LC601405.1 | | unidentified mushroom | fungus |
| *Kuraishia capsulata** | | Y72 | *Ips typographus*, this study | *Ips typographus* |
| *Kuraishia mediterranea* | PQ094279.1 | | Olive oil production process | food/fodder |
| *Kuraishia molischiana** | | Y88 | *Ips typographus*, this study | *Ips typographus* |
| *Nakazawaea ernobii* | KY108637.1 | | intracellular symbiont of *Ernobius mollis* | insect |
| *Nakazawaea holstii** | | Y141 | *Ips typographus*, this study | *Ips typographus* |
| *Nakazawaea wickerhamii* | KY108656.1 | | silage, containing olive husks, molasses and whey | food/fodder |
| *Ogataea cecidiorum* | KY108670.1 | | galls induced by sawflies on leaves of willows | insect |
| *Ogataea minuta* | KY106579.1 | | rotten flower of Rhododendron indicum | plant |
| *Ogataea pini* | KY108703.1 | | insect frass on *Pinus taeda* (loblolly pine) | insect |
| *Ogataea ramenticola* | KY108713.1 | | tunnel of insect in Pinus palustris (long-leaf pine tree) | insect |
| *Ogataea zsoltii* | KY108723.1 | | rotten railway-sleeper | decaying wood |
| *Saccharomyces bayanus* | MH318012.1 | | Riesling Icewine grapes from the Niagara Region | plant |
| *Saccharomyces boulardii* | OR786914.1 | | beer brewing process | food/fodder |
| *Sugiyamaella chuxiongensis* | NG228810.1 | | beer brewing process | decaying wood |
| *Sugiyamaella yunanensis* | NG228840.1 | | rotting wood | decaying wood |
| *Torulaspora indica* | KY109871.1 | | Coal mine soil, Singareni Collries | soil |
| *Torulaspora maleeae* | KY109872.1 | | from leaf Rhizophora mucronata | plant |
| *Trichomonascus apis* | NG055369.1 | | honeycomb | insect |
| *Trichomonascus ciferrii* | LC158141.1 | | human throat | human |
| *Wickerhamiella brachini* | NG079503.1 | | *Brachinus scotomedes* | insect |
| *Wickerhamiella fructicola* | NG59950.1 | | Fruits of white garland lily (*Hedychium coronarium*) | plant |
| *Wickerhamiella goesii* | NG243660.1 | | sugarcane leaves | plant |
| *Wickerhamomyces anomalus* | MT705623.1 | | *Rosa roxburghii* | plant |
| *Wickerhamomyces bisporus** | | Y143 | *Ips typographus*, this study | *Ips typographus* |
| *Wickerhamomyces menglaensis* | NG228761.1 | | rotting wood from different locations in Xishuangbanna | decaying wood |
| *Wickerhamomyces xylosivorus* | NG057186.1 | | Decayed wood | decaying wood |
| *Yamadazyma mexicana** | | Y5 | *Ips typographus*, this study | *Ips typographus* |
| *Yamadazyma paraaseri* | NG243041.1 | | rotting wood | decaying wood |
| *Yamadazyma scolyti** | | Y130 | *Ips typographus*, this study | *Ips typographus* |
| *Yamadazyma tenuis** | | Y63 | *Ips typographus*, this study | *Ips typographus* |

**Suppl. Table S2:** Overview of all fungal isolates used for the experiments in this study.

| Species | Accession | Isolate ID | Olfaction assay | Confrontation assays | Egg infection | Stilbene degradation | Growth on phenolic acids | Strip assays | Fungal biomass | Untargeted analysis |
|---|---|---|---|---|---|---|---|---|---|---|
| *Danielozyma ontarioensis* | this study | Y139 | x | | | | | | | |
| *Kuraishia capsulata* | this study | Y72 | x | x | x | x | x | | | x |
| *Kuraishia molischiana* | this study | Y88 | x | x | | x | x | | | |
| *Nakazawaea holstii* | this study | Y141 | x | x | | x | x | | | x |
| *Wickerhamomyces bisporus* | this study | Y143 | x | x | | x | x | | | x |
| *Yamadazyma scolyti* | this study | Y130 | x | x | | x | x | | | |
| *Yamadazyma tenuis* | this study | Y63 | x | x | | x | x | | | |
| *Trichoderma harzianum* | this study | P338 | x | x | x | | x | x | x | |
| *Grosmannia penicillata* | PQ897185 | NG481 | | x | | | | x | x | |
| *Grosmannia penicillata* | this study | 15fK2 | | | | | x | | | |
| *Ophiostoma bicolor* | PQ897178 | 460 | | | | | | | x | |
| *Endoconidiophora polonica* | PX884903 | P331 | | | | | x | | | |
| *Endoconidiophora polonica* | PQ897182 | NG471 | | x | | | | x | x | |
| *Ceratocystiopsis minuta* | PQ900763 | NG473 | | | | | x | | x | |
| *Graphium fimbriasporum* | this study | 4fW2 | | | | | x | | | |
| *Grosmannia europhioides* | this study | P333 | | | | | x | | | |
| *Chaetomium globosum* | MT252028 | P7 | | | | | x | | | |
| *Cylindrobasidium ipidiophilum* | this study | P334 | | | | | x | | | |
| *Ophiostoma floccosum* | this study | 6mK3 | | | | | x | | | |

### Supporting Text: *Kuraishia capsulata* metabolism

***Kuraishia capsulata* possesses a specialized stilbene metabolism.** While most *Ips typographus*-associated yeasts deglycosylate the spruce's stilbenes and thus produce antifungal stilbene aglycones, *K. capsulata* further breaks these compounds down into the respective phenolic acids (Fig. 5, Suppl. Figs. S4-5). To verify this and to investigate the mechanism of stilbene aglycone catabolism, we used resveratrol as a representative of the stilbene aglycones derived from spruce phenolic compounds. Liquid chromatography quadrupole time-of-flight mass spectrometry (LC-QTOF-MS) analysis showed that *K. capsulata* significantly reduced the levels of resveratrol in the growth medium compared to *W. bisporus* and the control (based on intensity of m/z 227.07083 from LC-Q-TOF-MS in negative mode, Suppl. Fig. S12a-b). We identified three compounds that were exclusively present in resveratrol-amended PDA inoculated with *K. capsulata*: (**1**) hydroxylated form of 3,5-dihydroxybenzoic acid, *m/z* = 169.01435; (**2**) 3,5-dihydroxybenzoic acid, *m/z* = 153.01943; and (**3**) 4-hydroxybenzoic acid, *m/z* = 137.0244 (Suppl. Fig. 9c). Further LC-QTOF-MS analysis of three representative yeasts grown in the presence of 3,5-dihydroxybenzoic acid showed that only *K. capsulata* depleted the growth medium of this phenolic acid (**2**) and it was the only isolate that produced a hydroxylated form of this compound (**1**) (Suppl Fig. S10). Our results point towards an oxidative cleavage of the double bond present in resveratrol, resulting in the production of 4-hydroxybenzoic acid and 3,5-dihydroxybenzoic acid which is further hydroxylated at a later step (Suppl Fig. S11).

### Supporting Text: Methods – analytical chemistry

Extraction

Methanol extracts were prepared from dried fungal biomass by homogenizing it with 1 mL of methanol (Honeywell) and three metal beads (Ø 3 mm, Askubal) on a paint shaker (Skandex SO-10M, Fluid Management Europe, The Netherlands) for 5min. Samples were centrifuged afterwards at 9400 rcf for 2 min and supernatant was collected for further analyses (1).

Analysis of stilbenes and phenolic acids by LC-MS/MS

Stilbenes were quantified with an LC-MS/MS system. We performed chromatography on an Agilent 1200 HPLC system (Agilent Technologies, Boeblingen, Germany), where the separation was achieved on an Agilent Zorbax Eclipse XDB-C18 column (50 x 4.6 mm, 1.8 µm; Agilent Technologies, Santa Clara, CA, USA). Formic acid (0.05%) and acetonitrile were employed in water as mobile phases A and B, respectively. The elution profile was: 0.0-0.5 min, 5% B; 0.5-6.0 min, 5-37.4% B; 6.0-6.02 min, 37.4-80% B; 6.02-7.5 min, 80-100% B; 7.5-9.5 min, 100% B and 9.5-12 min 5% B. The flow rate for the mobile phase was set to 1.1 mL/min while the column temperature was maintained at 25°C. The HPLC was coupled to an API 3200 tandem mass spectrometer (Applied Biosystems, Darmstadt, Germany) equipped with a Turbospray ion source, which was operated in the negative ionization mode. We optimized the instrument parameters by infusion experiments with pure standards. The ion spray voltage was maintained at -4500 eV and the turbo gas temperature was set at 500°C. Nebulizing gas was set at 60 psi, curtain gas at 30 psi, heating gas at 60 psi and collision gas at 4 psi. The mass spectrometer was operated in multiple reaction monitoring (MRM) mode, and details of the instrument parameters can be found in Suppl. Table S6. We maintained both Q1 and Q3 quadrupoles at unit resolution. Analyst 1.5 software (Applied Biosystems, Darmstadt, Germany) was used for data acquisition and processing. Target compounds were quantified based on external standard curves using dilution series of commercial standards (as listed in Suppl. Table S6). Samples were normalized by the individual sample dry weight unless fresh samples were extracted directly. In these cases, the fresh weight was used to normalize the concentrations.

**Suppl. Table S3.** Details of analysis of stilbenes and phenolic acids by LC-MS/MS [HPLC 1200 (Agilent Technologies)-API3200 (AB SCIEX)] in negative ionization mode

| Compound | Q1 | Q3 | RT (min) | DP | CE | Supplier |
|---|---|---|---|---|---|---|
| Protocatechuic acid | 153 | 108 | 2.7 | -35 | -28 | Santa Cruz Biotechnology, USA |
| Vanillic acid | 167 | 123 | 4 | -35 | -18 | Sigma Aldrich, Germany |
| 4-hydroxy-benzoic acid | 137 | 93 | 3.6 | -35 | -20 | Tokio Chemical Industry, Japan |
| Piceatannol | 242.9 | 159.1 | 5.3 | -55 | -34 | Tokio Chemical Industry, Japan |
| Isorhapontigenin | 256.9 | 240.9 | 6.4 | -35 | -28 | Tokio Chemical Industry, Japan |
| Resveratrol | 227 | 185 | 6.3 | -40 | -28 | Tokio Chemical Industry, Japan |
| Piceid | 389 | 227 | 5 | -50 | -38 | Tokio Chemical Industry, Japan |
| Astringin | 404.8 | 243 | 4.5 | -50 | -38 | Isolated by Ruo Sun |
| Isorhapontin | 418.9 | 257.1 | 4.9 | -25 | -18 | not available |

Untargeted metabolite analysis – Resveratrol metabolism in *Kuraishia capsulata*

To investigate the phenolic acid metabolism in *K. capsulata*, 100 µl of liquid yeast culture (*W. bisporus*, *K. molischiana*, OD600 = 0.1) were inoculated on PDA amended with either 2 % (v/v) DMSO or 200 µg/g resveratrol (n = 6 per yeast per treatment), all supplemented with a sterile cellophane disk (NeoLab, Germany) to facilitate sampling the agar without yeast biomass. The plates were incubated for 72 hours at 25°C and 65% relative humidity. A sample of agar was taken from each plate, freeze dried and extracted in methanol following the protocol described earlier. The extracts were used to perform untargeted metabolite analysis by LC-ESI-Q-ToF-MS.

To carry out untargeted metabolite analysis, ultra-high-performance liquid chromatography–electrospray ionization– high resolution mass spectrometry (UHPLC–ESI–HRMS) was performed with a Dionex Ultimate 3000 series UHPLC (Thermo Scientific) and a Bruker timsToF mass spectrometer (Bruker Daltonik, Bremen, Germany) as described in Müller *et al.* (2). UHPLC used a reversed-phase Zorbax Eclipse XDB-C18 column (100 mm × 2.1 mm, 1.8 µm, Agilent Technologies, Waldbronn, Germany) with a solvent system of 0.1% formic acid (A) and acetonitrile (B) at a flow rate of 0.3 ml/min. The elution profile was the following: 0 to 0.5 min, 5% B; 0.5 to 11.0 min, 5% to 60% B in A; 11.0 to 11.1 min, 60% to100% B, 11.1 to 12.0 min, 100% B and 12.1 to 15.0 min 5% B. Electrospray ionization (ESI) in negative ionization mode was used for the coupling of LC to MS. The mass spectrometer parameters were set as follows: capillary voltage 3.5 KV, end plate offset of 500 V, nebulizer pressure 2.8 bar, nitrogen at 280°C at a flow rate of 8 L/min as drying gas. Acquisition was achieved at 12 Hz with a mass range from *m*/*z* 50 to 1500, with data-dependent MS/MS and an active exclusion window of 0.1 min, a reconsideration threshold of 1.8-fold change, and an exclusion after 5 spectra. Fragmentation was triggered on an absolute threshold of 50 counts and acquired on the two most intense peaks with MS/MS spectra acquisition of 12 Hz. Collision energy was alternated between 20 and 50 V. At the beginning of each chromatographic analysis 10 µL of a sodium formate-isopropanol solution (10 mM solution of NaOH in 50/50 (v/v%) isopropanol water containing 0.2% formic acid) was injected into the dead volume of the sample injection for re-calibration of the mass spectrometer using the expected cluster ion *m*/*z* values.

The feature detection of LC-ESI-Q-ToF-MS raw data was carried out using Metaboscape software (Bruker Daltonik, Bremen, Germany) with the T-Rex 3D algorithm for qTOF data. For peak detection the following parameters were used: intensity threshold of 1000 with a minimum of 10 spectra, time window from 0.4 to 12 min, peaks were kept if they were detected in at least 60% of all replicates of one sample group. Adducts of $[M-H]^-$, $[M-H+Cl]^-$, and $[M-H+COOH]^-$ were grouped as a single bucket if they had an EIC correlation of 0.8.

The target compound (3,5-dihydroxybenzoic acid, marked as “peak 2” in the results) showed a signal at *m/z* of 153.0194 for the $[M-H]^{qq}$ ion ($C_7H_5O_4$, theoretical value: 153.0193; $\Delta$ -0.7 ppm) with a fragment ion at *m/z* 109.0293 ($C_6H_5O_2$) after collision-induced dissociation (CID) fully consistent with that of authentic 3,5-dihydroxybenzoic acid (Sigma Aldrich, Germany).

After identification of 3,5-dihydroxybenzoic acid as a unique breakdown product of resveratrol in the agar inoculated with *K. capsulata,* we repeated the cultivation experiment using this phenolic acid as a supplement to the growth media. In brief 100 µl of liquid yeast culture (*W. bisporus*, *N. holstii, K. molischiana*, OD600 = 0.1) were inoculated on PDA amended with 50 µg/g 3,5-dihydroxybenzoic acid (n = 5 per yeast per treatment) supplemented with a sterile cellophane disk (NeoLab, Germany). The plates were incubated for 5 days at 25°C and 65% relative humidity. A sample of agar was taken from each plate, freeze dried and extracted in methanol as described earlier. The extracts were used to perform untargeted metabolite analysis by LC-ESI-Q-ToF-MS as described above.
